# How demography shapes linkage disequilibrium with or without recombination

**DOI:** 10.1101/2023.12.13.571342

**Authors:** Elise Kerdoncuff, Mariadaria K Ianni-Ravn, Denis Roze, John Novembre, Amaury Lambert, Guillaume Achaz

## Abstract

Linkage Disequilibrium (LD) is a measure of the statistical association, within a population, between alleles present at different loci. Here, we produce detailed characterizations of LD statistics in different demographic scenarios, with simulations and with analytic results, focusing on several widely used measures of LD (*D*, |*D*′|, and *r*^2^), and introducing a fourth class of statistics: 2-site configuration probabilities, which correspond to the frequencies of different allelic association patterns. By studying these measures under no recombination, a single recombination event, and between fully independent loci, we are able to gain new insights to how these measures vary across recombination scales. We also classify recombination events into four types and rank them according to their respective frequencies and effects on LD measures. Using previously published human data, we show how tree-sequence information can be used to partition the genome and to analyze LD in relation to theoretical expectations. Finally, we provide recommendations for the use of the different statistics and suggest applications of LD measures in the absence of recombination.

## 1 Introduction

Linkage Disequilibrium (LD) quantifies the statistical association between variants located at two different loci (Lewontin and Kojima, 1960). At its appearance, a newly created variant (also known as a ‘derived allele’) is physically and therefore statistically associated with all other alleles located at all other loci of the genome in which it occurs. Due to the random segregation of chromosomes during meiosis, associations between alleles at loci belonging to different chromosomes will rapidly decrease, at a rate of 0.5 per generation. The statistical association between alleles at loci located on the same chromosome can last longer, as its decay is tuned by the recombination rate: the smaller the probability of recombination, the tighter the association and thus the longer the persistence time.

In the simple case of two biallelic loci, four haplotypes can segregate: *AB, Ab, aB*, and *ab*, where *A* and *a* are alleles at the first locus and *B* and *b* are alleles at the second locus. Let us assume *a* and *b* are the ancestral alleles, with *A* and *B* being derived. When a new mutation introduces the derived allele *A*, it can arise on either the *b* or *B* background. For example: if *A* arises on an *ab* haplotype, the initial mutant haplotype is *Ab*, while if *A* arises on an *aB* haplotype, the initial mutant haplotype is *AB*. Thus, at the time *A* appears, there are typically only three haplotypes present in the population, depending on the background on which *A* arises. For instance, if *A* arises on *ab*, the three haplotypes are *ab, aB*, and *Ab*; if it arises on *aB*, the haplotypes are *ab, aB*, and *AB*. More generally, in the absence of recombination (and recurrent mutation), a maximum of three of the four haplotypes segregate in the population, regardless of their frequency. The presence of the fourth haplotype is classically taken as a sign of past recombination events, a test known as the 4-gamete test (Hudson and Kaplan, 1985).

Let us denote *f*_*A*_ the frequency of *A* (*f*_*a*_ = 1 − *f*_*A*_), *f*_*B*_ the frequency of *B*, and *f*_*AB*_ the frequency of the haplotype *AB*. The canonical measure of LD is then defined as *D* = *f*_*AB*_ − *f*_*A*_*f*_*B*_ and originally framed as a measure of departure from an equilibrium (Lewontin and Kojima, 1960), but it’s also readily seen to be a covariance between allelic states of 2-loci on a haplotype. Further mathematical details, interpretations, and normalizations of *D* are provided below in the Methods section.

As LD is affected by virtually all evolutionary and population-genetic processes, it can be used to infer forces as diverse as recombination (McVean *et al*., 2004), natural selection (Sabeti *et al*., 2002), demography (Li and Durbin, 2011; Harris and Nielsen, 2013; Ragsdale and Gravel, 2019), population structure (Pritchard *et al*., 2000), founder events (Tournebize *et al*., 2022), and gene–trait associations (Visscher *et al*., 2017; Uffelmann *et al*., 2021) (see also reviews by Pritchard and Przeworski (2001); Nordborg and Tavaré (2002); Slatkin (2008); Balding *et al*. (2008)). We will first briefly review the effect of mutation and recombination (intrinsic genetic factors) before assessing the effect of genetic drift and selection (population processes).

### Mutation and recombination

As mentioned above, new pairs of alleles initially arise in complete association, producing genealogical LD driven by their shared local ancestry. As a result, newly created variants segregating at low frequency typically show strong associations with neighboring variants.

On the contrary, recombination events shuffle existing variants located at the two loci and break the statistical association between loci. The magnitude of LD depends on the number of past recombination events between the two loci (Lewontin and Kojima, 1960), that itself depends on the genetic distance, that integrates recombination rate and physical distance (Long and Langley, 1999). In humans, the average per-base recombination probability is roughly 10^−8^ per generation. For variants within a typical exon of 200bp, the per-generation recombination probability is only 2 × 10^−6^, implying that LD between such closely linked sites can persist over many generations before being broken by recombination. As a result, the expected value of *D* decreases geometrically over *t* generations at a rate *c*,

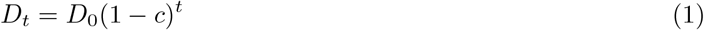

where *D*_0_ is an initial value of *D* and *D*_*t*_ is the value after *t* time steps, and where *c* is the per generation probability of having an odd number of crossovers between two sites (if there is an even number of crossovers between two sites, they end up on the same gamete). This geometric decay corresponds to an exponential decay *D*_*t*_ ≈ *D*_0_*e*^−*ct*^ for small *c*, which is typically the regime of interest for closely linked sites. If *r* denotes the recombination probability per base pair and *L* the physical distance between the two sites, then in the absence of interference between crossovers: *c* = (1 − *e*^−2*rL*^)*/*2 ≈ *rL* for *rL* ≪ 1 (Haldane, 1919).

### Genetic drift

The strength of LD is not only determined by recombination but also by population size. In finite populations, stochastic fluctuations in allele frequencies (genetic drift) generate non-random associations between loci. Importantly, the magnitude of these associations is inversely related to the effective population size, such that smaller populations tend to exhibit higher levels of LD.

Under this framework, the sampling of alleles in each generation can create LD between a pair of loci even if they were previously at equilibrium. However, for a given pair, the expected value of *D* converges to zero over time. Early work further showed that, in the absence of selection, the mean of *D across pairs* is zero, while its variance depends on population size and the frequency of recombination. Hill and Robertson (1968) obtained an exact analytical solution for the case of no recombination and no selection using a moment generating matrix, whereas Ohta and Kimura (1969b) derived the variance of *D* across pairs in the presence of recombination using Kolmogorov backward equations. Classical two-locus diffusion theory indicates that, in the limit of large effective population size *N* and small recombination rate *c*, the variance of LD generated by drift scales inversely with the population-scaled recombination parameter *Nc*. In particular, many summaries of the stationary two-locus solutions imply the leading behavior

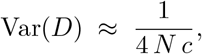

highlighting that LD due to drift is stronger in smaller populations or between more tightly linked loci (see, e.g., Hill and Robertson 1968; Ohta and Kimura 1969b; McVean 2002).

It is important to note that these expectations of the variance of *D* include cases in which one or both loci are monomorphic (when *D* is necessarily 0) (Slatkin, 2008). The variance of *D* when both loci are polymorphic has been derived analytically for multiallelic loci (Hill, 1974; Hill and Weir, 1988), showing that it depends on population size and allele frequencies. Genetic drift continuously generates LD, while recombination erodes it, resulting in a background level of LD maintained by mutation–drift–recombination dynamics, especially between loci separated by only a few cM on the same chromosome. As background LD is expected to be strong at short distances, it is often suggested closely located markers should be thinned before carrying out analyses that model LD generated only by other sources, such as admixture LD (Falush *et al*., 2003); although the exact distance concerned is not clear and will depend on the species studied, and for some applications, such as detecting inbreeding (Weir and Hill, 1980; Golding and Strobeck, 1980) or recent positive selection, removing close markers may reduce power.

### Selection at linked neutral sites

Natural selection causes beneficial variants to increase in frequency. When the time scale of fixation is short, recombination has few opportunities to break associations between the selected variant and nearby alleles. As a result, these surrounding alleles are carried along to high frequency, a phenomenon known as hitchhiking (Smith and Haigh, 1974). This process creates strong positive LD (*D >* 0) between neighboring alleles found on the ancestral haplotype of the selected allele, a signal that can be used to detect ongoing or past selective sweeps (Sabeti *et al*., 2002). Background selection (BGS) can have a similar but more diffuse effect. By continually removing deleterious mutations, BGS reduces the effective population size, increasing the influence of genetic drift (Charlesworth, 1994; Hudson and Kaplan, 1995). Since LD scales inversely with effective population size, BGS is expected to elevate baseline LD across linked loci, even in the absence of positive selection.

### Selection between loci

Selection can also generate LD through interactions between loci, such as epistasis. Positive LD (*D >* 0) indicates that alleles co-occur more often than expected under independence, as may happen with synergistic interactions between beneficial alleles (Lewontin and Kojima, 1960). Negative LD (*D <* 0) suggests that alleles are more often found on different haplotypes, which can arise due to interference between deleterious mutations, a phenomenon known as Hill-Robertson interference (Hill and Robertson, 1966) or due to stabilizing selection on a quantitative trait, a phenomenon known as the Bulmer effect (Bulmer, 1971). Patterns of signed LD between selected loci can thus provide insights into the nature of the selective forces acting across the genome, including epistatic interactions.

### Other evolutionary influences

LD reflects the full evolutionary history of the observed alleles, *i*.*e*. the population history, the breeding system as well as the pattern of geographic subdivision (Slatkin, 2008). For example, mixture or admixture of populations with different ancestries generate patterns of LD, with alleles from the same ancestry in strong positive LD and alleles from different ancestries in strong negative LD (Ohta, 1982). Inbreeding generates another pattern, with long continuous blocks of alleles in strong LD as a result of being inherited from the same common ancestor (that is, blocks inherited identical by descent) (Charlesworth, 2003). Gene conversion produces similar patterns as recombination; it breaks down LD between alleles (even in the case of homologous gene conversion as some fraction of present-day chromosomes will show some recent allele surrounded by ancestral background) (Jones and Wakeley, 2008). Therefore, LD can be studied in bacterial populations (Haubold *et al*., 1998) and, for example, be used to detect selection (Rocha, 2018).

When analyzing LD empirically, one can use it to infer evolutionary history and population-genetic processes. First, it is common to study how LD decays with genetic distance (on average exponentially, even once time is integrated in eq. 1). To assess it, measures of LD are viewed as functions of genetic distance. As the rate of this decay varies with processes such as demography, it can be used to estimate demographic parameters (Loh *et al*., 2013; Boitard *et al*., 2016). Second, patterns of LD along the genome and particularly the length of segments in LD are often used for inference. The smaller the population is, the more individuals share segments in LD, which makes haplotype/segment length a good indicator of past population sizes (Schiffels and Durbin, 2014; Kerdoncuff *et al*., 2020; Tian *et al*., 2019). In addition, extremely long haplotypes shared among a large fraction of the sample can be a sign of past positive selection events (Sabeti *et al*., 2002; Voight *et al*., 2006).

Interpretation of LD can be challenging because most analytic results are derived under simplified assumptions, such as neutrality or constant population size. Although theoretical work has developed approaches to address biases and more complex scenarios, in practice it remains difficult to account for multiple evolutionary processes simultaneously. For example, measures of signed LD can be biased when focusing on rare alleles, highlighting the importance of considering allele frequency and sampling effects (Ragsdale, 2022; Good, 2022). This complexity has motivated the development of more sophisticated LD-based metrics to disentangle the contributions of different processes (Kaeuffer *et al*., 2007).

As described, LD-based methods often need genetic distance, however, turning physical distance into genetic distance requires prior knowledge of recombination rate and/or the genetic map. Thus, the decay of LD with physical distance can be used to estimate the recombination rate (Pritchard and Przeworski, 2001; McVean *et al*., 2002; Auton and McVean, 2007; Chan *et al*., 2012). One complication arises as, within an individual, the recombination rate varies along the genome (Kong *et al*., 2002; McVean *et al*., 2004). These local variations of the recombination rate are a major source of modulation in LD values (Altshuler *et al*., 2005). The use of genetic distance (when not inferred using LD) instead of physical distance results in LD values that are independent of these local variations. The inference of genetic maps nonetheless is very time consuming and necessitates either controlled crosses (Campbell *et al*., 2016) or good knowledge of the population pedigree (Kong *et al*., 2002). Recombination rates also vary across species (Smukowski and Noor, 2011; Hewett *et al*., 2023), across populations and between sexes (Kong *et al*., 2010). Therefore, fine-scale recombination maps are difficult to obtain, even for a given species. In addition, the large majority of population genetic methods to estimate recombination rates assume simple neutral demographic models (see for a notable exception pyrho (Spence and Song, 2019) which allows for step-wise changes in population size).

In practice, LD measures in the absence of recombination events can be estimated between pairs of sites that are sufficiently close on the genome such that no recombination is expected to have occurred between them. When the sequence alignment can be divided into maximal recombination-free blocks (Kerdoncuff *et al*., 2020), these statistics simply reflect the LD patterns observed within each block. More broadly, recent advancements in reconstructing ancestral recombination graphs (ARGs) and encoding genealogical histories as tree sequences offer a powerful framework for pinpointing the precise locations and numbers of recombination events across the genome (Rasmussen *et al*., 2014; Kelleher *et al*., 2019; Speidel *et al*., 2019; Deng *et al*., 2025). Within a tree sequence, the span of a single tree naturally defines a scale over which genealogical LD is expected to predominate, offering an intuitive connection between genealogical structure and observed statistical associations. By leveraging these representations, it becomes possible to condition LD statistics on the number of recombination events between loci. In this context, we also investigate the effect of a single recombination event on LD patterns. This will allow us to explore the consequences of failing to detect a recombination event and instead calculating LD measures as if no recombination had occurred between loci. For comparison, we also examine the LD measures under the assumption of infinite recombination, where loci are considered independent.

In this article, we seek to systematically study the effect of recombination and demography on LD, focusing mostly on three established statistics: *D*, |*D*′| and *r*^2^ along with newly defined metrics called 2-site Configuration Probabilities (CPs). We study patterns of LD for these statistics, in three different cases: absence of recombination, one single recombination event and an infinite number of events. Demographic changes are modeled as a single instantaneous population size change. As a proof of concept, we measure these statistics on previously inferred tree sequences from the 1000 Genomes Project, demonstrating their potential for evolutionary inference.

## 2 Materials and Methods

As an overview, we analyze various metrics of Linkage Disequilibrium, namely, *D*, |*D*′|, *r*, 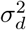 and *r*^2^ and introduce a new set of metrics called CPs. For most of them, we estimate their distributions (often only the means and variances) using Monte-Carlo simulations, varying the number of recombination events (0, 1 or an infinity) and the demography of the population using a 2-epoch model. As an application, we also compute these LD metrics in human population data.

### 2.1 Metrics considered

LD can be quantified using a variety of metrics, which we classify here into pairwise measures for individual locus pairs and aggregate measures that summarize patterns across many pairs of loci.

#### Pairwise metrics

These metrics quantify the statistical association between a specific pair of alleles:

- *D* = *D*_*AB*_ = *f*_*AB*_ − *f*_*A*_*f*_*B*_ (Lewontin and Kojima, 1960). *D* is a covariance and consequently signed. *D* decays geometrically (eq. 1, and approximately exponentially for small *c*) with the number of recombination events between both loci (both are proportional to *t* and *c*). “Linkage equilibrium” is when *D* = 0.
- |*D*′| = |*D*| */* max(|*D*|), where max(|*D*|) is the maximal value that |*D*| can take when *f*_*A*_, *f*_*B*_, and the sign of *D* are known and fixed, but *f*_*AB*_ can vary (Lewontin, 1964). It can be shown that knowing *f*_*A*_, *f*_*B*_ and the sign of *D*, |*D*| is maximal whenever one the four combinations of alleles is missing and is then equal to min(*f*_*A*_*f*_*b*_, *f*_*a*_*f*_*B*_) if *D >* 0 and to min(*f*_*A*_*f*_*B*_, *f*_*a*_*f*_*b*_) if *D <* 0. In particular, |*D*′| deviates from 1 only when the four combinations of alleles are observed. It is known that |*D*′| varies linearly with *f*_*AB*_ (Kang and Rosenberg, 2019).
- 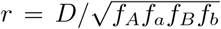. *r* is the Pearson coefficient of correlation between the presence of the two alleles. The covariance *D* is simply normalized by the standard deviations of the random variables, 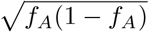 and 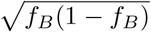, so that *r* ranges between −1 and 1.
- *r*^2^ = *D*^2^*/*(*f*_*A*_*f*_*a*_*f*_*B*_*f*_*b*_) (Hill and Robertson, 1968). The coefficient of determination (the square of *r*, the Pearson coefficient of correlation), varies quadratically with *f*_*AB*_ (Kang and Rosenberg, 2019).

#### Aggregate metrics

These summarize LD patterns across multiple pairs of loci rather than a single locus pair:

- 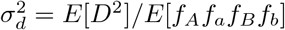 (Ohta and Kimura, 1971) is similar to *r*^2^, but uses the ratio of expectations of *D*^2^ and *f*_*A*_*f*_*a*_*f*_*B*_*f*_*b*_ and rather than the expectation of the ratio. Since 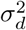 behaves similarly to *r*^2^ in most population-genetic contexts, we do not include it in all simulations and mention it primarily for comparison.
- CPs quantify the relative frequencies of different types of LD across a sample. For each pair of polymorphic sites, the correlation *r* between the derived alleles falls into one of four categories:
  – Perfect positive LD (*r* = 1): the derived alleles occur in exactly the same individuals.
  – Partial positive LD (0 *< r <* 1): the derived alleles co-occur more often than expected under independence, but not perfectly.
  – Partial negative LD (−1 *< r <* 0): the derived alleles co-occur less often than expected, but not perfectly opposite.
  – Perfect negative LD (*r* = −1): the derived alleles occur in completely non-overlapping sets of individuals.

CPs are then defined as the frequencies of pairs of sites in each of these four categories across the sample: {CP_Perfect Positive_, CP_Partial Positive_, CP_Partial Negative_, CP_Perfect Negative_}. By construction, the four probabilities sum to 1. CPs complement classical LD metrics by capturing the distribution of allele configurations rather than just covariances or correlations.

Independence between the alleles at the two loci results in a joint frequency equal to the product of the individual frequencies and consequently zero LD: *D* = |*D*′| = *r* = *r*^2^ = 0. How the values of the four statistics change in the presence of linkage (no independence) can greatly vary (McVean, 2007; Kang and Rosenberg, 2019). As an illustration, let us focus on the case when *A* and *B* are both singletons: *f*_*A*_ = *f*_*B*_ = 1*/n* for a sample of size *n*. Then we have: 1) *D* = 1*/n* − 1*/n*^2^ if *AB* exists and *D* = −1*/n*^2^ otherwise (in both cases *D* ≈ 0 for large *n*), 2) |*D*′| = 1 as at least one haplotype is missing, 3) *r* = 1 if *AB* exists and *r* = −1*/*(*n* − 1) ≈ 0 for large *n* if *AB* doesn’t exist, 4) *r*^2^ = 1 if *AB* exists and *r*^2^ = 1*/*(*n* − 1)^2^ ≈ 0 for large *n* if *AB* doesn’t exist and 5) a Perfect Positive configuration if *AB* exists and a Partial Negative configuration if *AB* doesn’t exist.

In this study, to assess the impact of recombination and evaluate the relevance of analyzing LD in its absence, we consider the different statistics conditional on the number of recombination events *k* that occurred between the two loci since their most recent common ancestor. The extended metrics are noted: 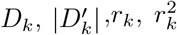 and CP_*k*_. We will focus for this study on three cases: *k* = 0 (no recombination), *k* = 1 (a single recombination event) and *k* → ∞ (independent loci).

### 2.2 Modeling assumptions

We now introduce the modeling assumptions and procedures by gradually adding the three layers of stochasticity: genealogies, recombination, and mutations.

#### 2.2.1 Genealogies and recombination

The genetic ancestry of sampled sequences is described by a sequence of local genealogies along the genome. In the absence of recombination, or when recombination is either absent or extremely rare, the data are described by a single tree. In contrast, free recombination leads to independent genealogies across loci.

We modeled recombination according to the ARG (Griffiths and Marjoram, 1997). Along the genome, recombination breakpoints occur as a Poisson process with rate *ρ*, the scaled recombination rate. More precisely, if *L* = *λ*(*T*_*k*_) denotes the total branch length of the local genealogy *T*_*k*_, then the distance to the next recombination breakpoint is exponentially distributed with rate *ρL*. Conditional on *T*_*k*_, a recombination event is placed uniformly along the tree: the probability that it occurs on a given branch *e* is proportional to its length, *L*(*e*)*/L*, analogous to the placement of mutations in the coalescent. Consequently, trees with greater total branch length are more likely to experience recombination over shorter genomic intervals. Furthermore, within a given tree, the timing of a recombination event along the genealogy depends on the number of lineages present at each moment: recombination is more likely to occur during periods when many ancestral lineages coexist.

At the recombination time, the lineage carrying the breakpoint is detached from the tree at the recombination time and subsequently re-coalesces according to Kingman’s coalescent dynamics. We retain all recombination events, including those for which the detached lineage re-coalesces with the same ancestral branch and thus leaves the local genealogy unchanged. When the re-coalescence occurs on a different branch, a new local genealogy *T*_*k*+1_ is obtained; otherwise, the genealogy remains identical to *T*_*k*_. This procedure generates a sequence of local trees along the genome, possibly with consecutive segments sharing the same topology and branch lengths.

In the special case where at most one recombination event occurs in the sample history, the SMC′(Marjoram and Wall, 2006) process coincides with the corresponding restriction of the full ARG, providing an exact description of the genealogical process used in our simulations.

#### 2.2.2 Mutations under the infinite-sites model

We assume the infinite-sites model, so that each polymorphic site is bi-allelic. Rather than explicitly simulating mutation events, we compute expectations over possible mutation events by considering the genealogy *T* at a given site and using the branch lengths to determine the probability that a mutation occurs on each edge. If *L* = *λ*(*T*) denotes the total length of the tree *T* (up to the most recent common ancestor in *T*) and *L*(*e*) denotes the length of edge *e*, then the probability of observing a derived allele at the tips of the subtree of descendants of *e* is the relative length of *e, L*(*e*)*/L*. The same procedure applies to any pair of polymorphic sites, independently conditional on their genealogies.

In empirical datasets, ancestral states are often uncertain. In such cases, we replace allele frequencies with minor allele frequencies (MAF), defined as 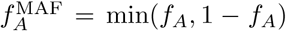 for locus 1 and analogously for locus 2. This ensures that analyses remain consistent when the derived state cannot be reliably identified.

### 2.3 Monte-Carlo simulations

All simulations in the main text were performed with a dedicated coalescent simulator (see Data/Software availability). For each scenario, we consider two coalescent trees, one for each locus assuming a sample of *n* = 10 haploid individuals (unless specified otherwise). Without recombination (*k* = 0), the tree is the same for both loci. For one recombination event (*k* = 1), the two trees are given by a simple ARG (Griffiths and Marjoram, 1997). With two loci and one recombination event, only one recombination event needs to be coded as in Marjoram and Wall (2006). For independent loci (*k* → ∞), the two trees are generated independently.

In Kingman simulations, each tree is sampled independently and with equal probability. In contrast, trees along the genome follow a different distribution: trees are generated sequentially along the genome, inducing correlations between adjacent trees. When a tall tree occurs, another tall tree is more likely to follow, even though tall trees typically span shorter genomic segments. If one samples a base pair uniformly along the genome, the local tree at that position follows the Kingman coalescent distribution. In this case, trees are effectively sampled in proportion to the length of genome over which they persist. However, sampling trees uniformly, ignoring their span, induces a different distribution, corresponding to a length-biased version of the Kingman law. This discrepancy implies that simulations based on independent trees do not directly reproduce the distribution relevant for mutation-based statistics. To correct for this, a weighting scheme is required so that each tree contributes proportionally to its genomic span, thereby recovering the appropriate distribution of trees seen by mutations (Kerdoncuff *et al*., In prep).

We do not generate polymorphic sites but instead take advantage of the branch lengths of the two trees to compute the probability of each possible sample configuration at the two sites, conditional on the two sites being polymorphic. This yields the *expected LD conditional on the genealogy(s)*, free from the mutational noise present in observed LD, which is exact in simulations where the underlying trees are known.

This procedure can be generalized to a two-site configuration, by sampling the pair of trees at each site and throwing uniformly at random one mutation over each of the trees, independently.

The distributions, the means and the variances are computed on 10^4^ replicates for a given scenario.

#### 2.3.1 Demography

We have implemented a simple 2-epoch demographic scenario. The population is modeled to be of size *N*_0_ at the time of sampling. Time is counted backward in units of *N*_0_ generations (coalescent time scale). We assume a single abrupt change of population size at time *τ* = 0.5 of strength *κ* = *N*_∞_*/N*_0_. Therefore, *κ* = 1 means no change, *κ >* 1 is a sudden decline and *κ <* 1 is a sudden growth (Figure 2).

### 2.4 Data analysis

To assess patterns of LD and evaluate the robustness of our predictions, we analyzed the 1000 Genomes Phase 3 (1KG3) dataset (The 1000 Genomes Project Consortium, 2015), which provides phased genotype data from 27 globally distributed human populations. Using previously inferred tree sequences inferred with tsinfer (Kelleher *et al*., 2019), we sub-sampled, for each population, ten independent sets of ten haploid genomes. The two haplotypes from each diploid individual were assigned to different sets to reduce diploid sampling correlations and approximate a haploid sampling scheme. The LD statistics discussed above were computed from phased variant calls in VCF files, averaging over observed pairs of biallelic sites restricted to the genomic span of each inferred tree. LD was evaluated at three recombination scales: zero recombination (within a tree), one recombination (across successive trees), and effectively infinite recombination (trees on different chromosomes). Trees with fewer than three sites or spans exceeding 10^5^ base pairs were excluded from the analysis. For distances 0 and 1, we sampled every hundredth tree across all autosomes; for distance ∞, we randomly sampled 100 tree pairs drawn from different chromosomes.

## 3 Results

### 3.1 No recombination

We first focus on cases when no recombination occurred between the two loci. The covariance between two variants with no recombination corresponds to two mutations occurring in the same coalescent tree.

#### 3.1.1 Analytical development

Analytical results for LD can be obtained using classical moment-based approaches, which derive expectations and variances of *D* by averaging over the joint distribution of genealogies under recombination (e.g. Hill and Robertson 1968; Ohta and Kimura 1969b; McVean 2002). In addition to these approaches, we develop in Supplementary Information I (6) a complementary framework that computes LD moments conditional on a given genealogy.

This genealogy-based approach provides exact expressions for the mean and variance of *D*_0_ for any fixed tree, without requiring averaging over the coalescent process. It is therefore particularly well suited to settings where genealogies are known or inferred (e.g. from tree sequences), and applies to arbitrary trees, whether ultrametric or not.

As specified earlier, any bi-allelic site has undergone a unique mutation. The frequency of the mutation in the sample conditional on the tree is random, assuming that the mutation falls on edge *e* with probability equal to the relative length of *e*, equal to *L*(*e*)*/L*, where *L*(*e*) is the length of edge *e* and *L* is the total length of the tree. As a consequence, the expectation *q* conditional on the tree, of the frequency *f*_*A*_ of the derived allele is

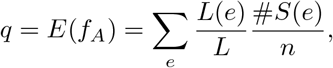

where *n* is the sample size, #*S*(*e*) is the number of descendants of edge *e* and the sum is taken over all edges of the tree. It is easily shown that

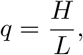

where we recall that *H* is the depth of the tree. Similarly, the expectation of the frequency *f*_*AB*_ of the double derived haplotype is *H*^2^*/L*^2^, so that

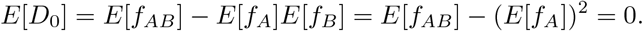

More involved calculations (Supplementary Note I (6)) show that

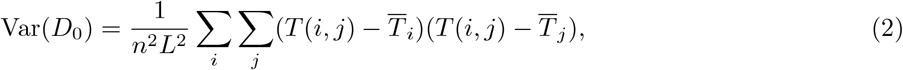

where each sum is taken over all tips of the tree, *T* (*i, j*) is the coalescence time between tip *i* and tip *j* and 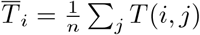.

Equation (2) shows that the variance of *D*_0_ is entirely determined by the covariance structure of pairwise coalescent times within a single genealogy. Importantly, this result holds conditional on a fixed tree, and therefore applies to any realization of the genealogy, including non-ultrametric trees such as those obtained from inferred ancestral recombination graphs or from data with heterogeneous sampling times.

This perspective complements the framework of McVean (2002), where LD is expressed in terms of covariances between coalescent times at two loci, averaged over the distribution of genealogies under recombination. In the limit of no recombination, both loci share the same genealogy, and McVean’s expression reduces to a combination of covariances between pairwise coalescent times within a single tree. In this case, it is algebraically equivalent to Eq. (2).

Our result can thus be interpreted as the single-genealogy counterpart of this broader coalescent framework, providing an exact expression for LD variance at the level of an individual tree.

This approach can be directly applied and verified in simulations: for each simulated tree, Eq. (2) provides the expected variance of *D*_0_, enabling comparison between analytical predictions and Monte-Carlo distributions obtained from placing mutations on the tree (see next section).

When studying the distribution of LD measures, we omit plots for *r*-based statistics (*E*[*r*_*k*_] and *Var*(*r*_*k*_)) because they are strongly correlated with other metrics. While *D*_0_ has mean zero (*E*[*D*_0_] = 0), the expectation of 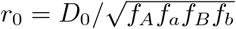 is not exactly zero, since the expectation of a ratio differs from the ratio of expectations. Nonetheless, *r*_0_ is symmetrically distributed around zero for typical sample sizes, so 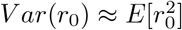 and analyzing *E*[*D*_0_] and 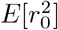 captures the main behavior of *r*_0_.

#### 3.1.2 Tree topology and allele frequencies

*D* is computed from allele frequencies at the two loci. Although the normalization of |*D*′| is still sensitive to allele frequencies, *r* and *r*^2^ were designed so that the output value is less sensitive to the frequencies. For example, when the two variants are systematically carried by the same haplotype (say only *AB* and *ab* are observed), *r* = *r*^2^ = 1 regardless of the frequency of the haplotype, contrary to *D* whose value will differ depending on *f*_*AB*_.

To illustrate this, we have computed in Figure 1, *D*_0_, *r*_0_ and 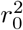 as a function of the frequency of *f*_*A*_ and *f*_*B*_. We consider two cases where the two mutations belong to the same subtree, called ‘nested’ *D*^(*n*)^; *r*^(*n*)^; *r*^2(*n*)^ and when they happened on different subtrees, called ‘disjoint’ *D*^(*d*)^; *r*^(*d*)^; *r*^2(*d*)^. These configurations lead to positively and negatively associated alleles, respectively.

**Figure 1:**
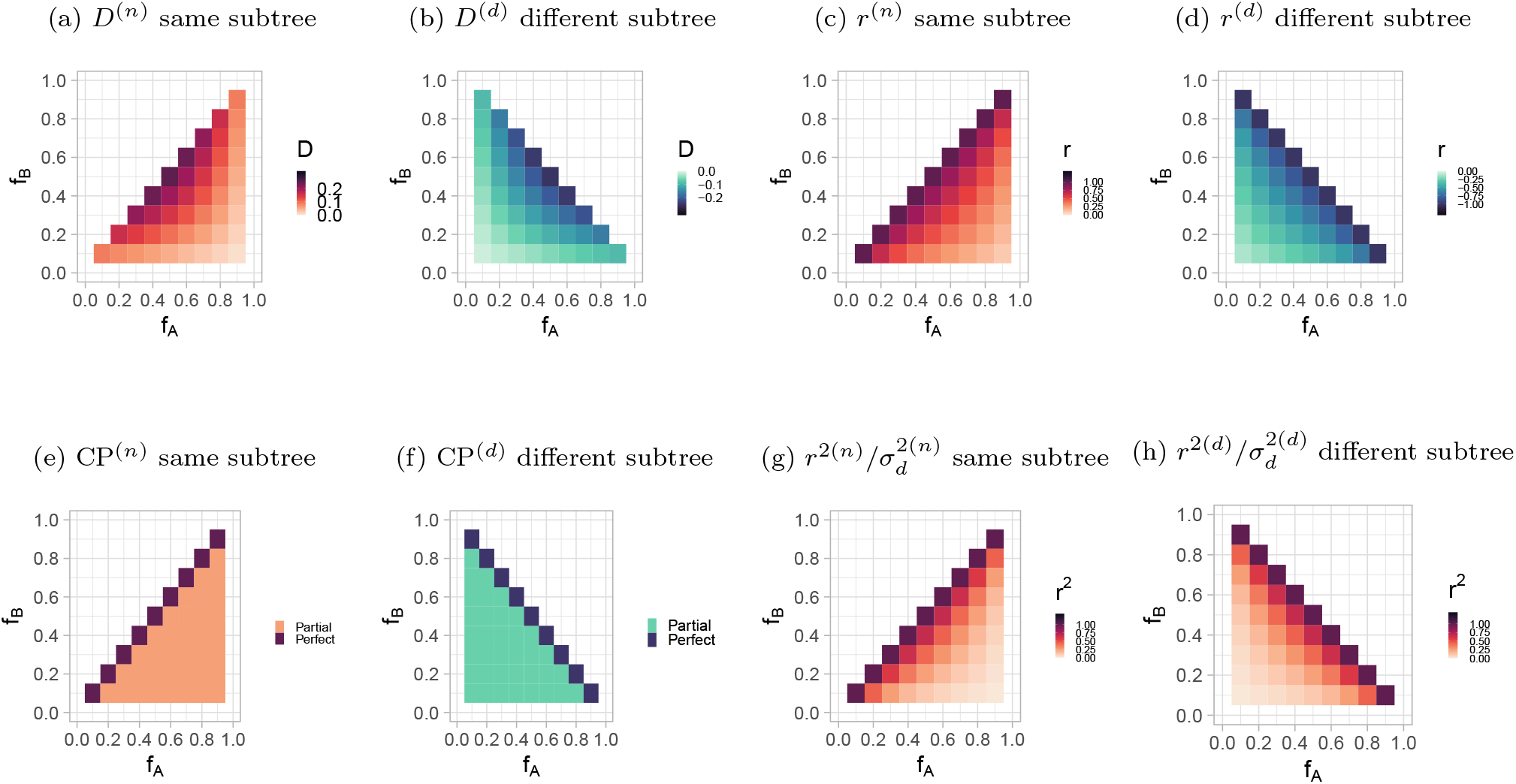
LD measures *D* (Figure 1a,1b), *r* (Figure 1c,1d), CP (Figure 1e,1f) and *r*^2^ (Figure 1g,1h) of two alleles A and B on the same tree, in function of their frequency *f*_*A*_ (x-axis) and *f*_*B*_ (y-axis). Two different scenarios, they happened on the same subtree ‘nested’ (left column) or on different subtree ‘disjoint’ (right column).

**Figure 2:**
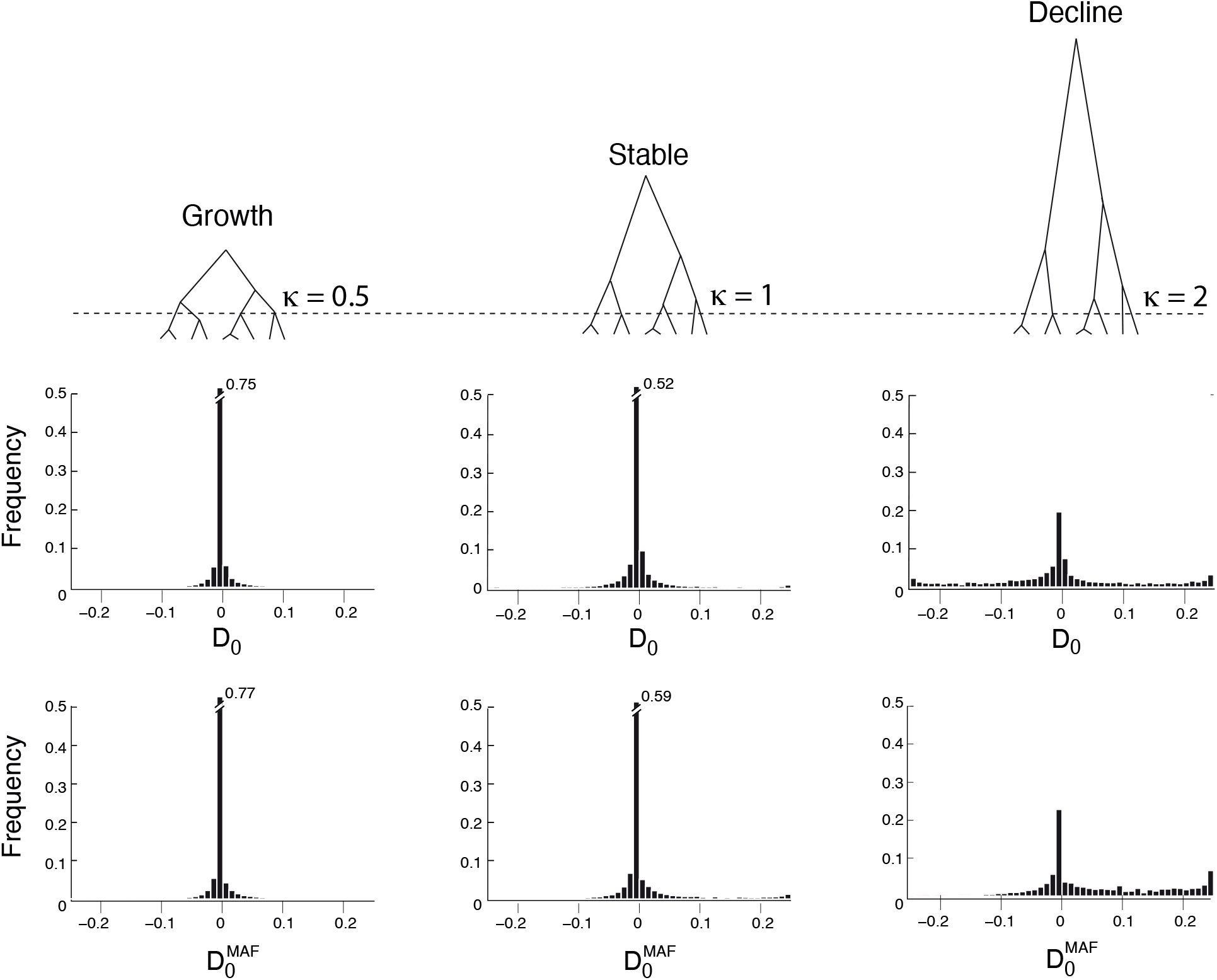
Distribution of *D*_0_ assuming *A* and *B* are the derived alleles (top panels) and *D*_0_ computed on the MAF 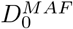 (bottom panels) under three demographic scenarios illustrated above the top panels: “Growth” is *κ* = 0.1, *τ* = 0.5, “Stable” is *κ* = 1, “Decline” is *κ* = 10, *τ* = 0.5. Sample size is *n* = 100 individuals and 1,000 trees were simulated per scenario.

*D*_0_: If alleles are nested with *f*_*A*_ ≥ *f*_*B*_: *f*_*B*_ = *f*_*AB*_; *D*_*AB*_ = *f*_*B*_(1 − *f*_*A*_), *D*^(*n*)^ values are positive (Figure 1a), whereas if they are disjoint: *f*_*AB*_ = 0; *D*_*AB*_ = −*f*_*A*_*f*_*B*_, *D*^(*d*)^ values are negative (Figure 1b). The extreme values of *D* are observed for medium allele frequencies. Alleles with low frequencies, as singletons, have *D* close to zero.

In the absence of recombination, the distribution of D values for a given tree can be computed using the two-Sites Frequency Spectrum (or 2-SFS), i.e the density of pairs of sites with mutation frequencies at *f*_*A*_ and *f*_*B*_, denoted *ξ*(*f*_*A*_, *f*_*B*_). Without recombination, 2-SFS can be divided into two different components: one nested component *ξ*^(*n*)^ for cases where the two mutations happened in the same subtree and some individuals carried the two mutations, and a disjoint component *ξ*^(*d*)^ that includes disjoint mutations that are only present in different individuals (Ferretti *et al*., 2018) and so, *ξ*(*f*_*A*_, *f*_*B*_) = *ξ*^(*n*)^(*f*_*A*_, *f*_*B*_) + *ξ*^(*d*)^(*f*_*A*_, *f*_*B*_). To obtain the distribution of D values we simply need to weigh the density of each pair *f*_*A*_, *f*_*B*_ by the values of *D*: *D*_*AB*_ = *D*^(*n*)^*ξ*^(*n*)^(*f*_*A*_, *f*_*B*_) + *D*^(*d*)^*ξ*^(*d*)^(*f*_*A*_, *f*_*B*_).

*r*_0_: Interestingly, the magnitude of *r* does not vary with absolute allele frequencies, but rather with the difference between them. Its sign will be the same as *D* but its absolute value will change. In a nested tree, *D* is positive and *r* is maximised when the alleles have the same frequency (*f*_*A*_ = *f*_*B*_). In a disjoint tree, *D* is negative and *r* is maximised when *f*_*A*_ = 1 − *f*_*B*_.

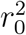 **and** 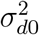: The range of values for *r*^2^ (equivalently 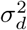 in this context) is not affected by the location of mutations. If mutations happened in the same subtree or in two different subtrees, the observed values can be similar (Figure 1g, Figure 1h), with a maximal *r*^2^ = 1 when *f*_*A*_ = *f*_*B*_ and *f*_*A*_ = 1 − *f*_*B*_. Two singletons on the same branches have a *r*^2^ = 1. Using the normalized LD-measure *r*^2^, we remove the allele frequency information which can be useful to understand the structure of the tree: the existence of distinct subtree or not and the weight of those subtrees if they exist.

**CP**_0_**s:** For a single genealogy, the different configurations can be directly related to the joint allele frequencies at the two sites: nested branches correspond to *f*_*AB*_ *>* 0 (positive associations), while disjoint branches correspond to *f*_*AB*_ = 0 (negative associations). The probabilities of these arrangements depend on branch lengths and the overall topology of the genealogy. Using allele frequencies, the four types of CP_0_ are defined as follows:

- **Perfect positive:** Both mutations occur on the same branch (*f*_*AB*_ = *f*_*A*_ = *f*_*B*_ = *f*; *D* = *f* (1 − *f*), *r* = 1).
- **Partial positive:** Mutations occur on branches within the same subtree, with one nested within the other but not on the same branch (*f*_*A*_ *> f*_*B*_, *f*_*B*_ = *f*_*AB*_; *D* = *f*_*B*_(1 − *f*_*A*_), 0 ≤ *r <* 1).
- **Perfect negative:** Mutations occur on completely opposite branches descending from the root (*f*_*A*_ = 1 − *f*_*B*_; *D* = −*f*_*A*_(1 − *f*_*A*_), *r* = −1).
- **Partial negative:** Mutations occur on different disjoint branches within the tree (*f*_*A*_≠ 1 − *f*_*B*_; *D* = −*f*_*A*_*f*_*B*_, −1 *< r* ≤ 0).

Simulations under a neutral coalescent model reveal that two mutations are more likely to be in a negative configuration (Figure 3 for *κ* = 1) than in a positive one, suggesting a “background” of subtle negative LD. However, this does not imply that the mean *D*_0_ should be different from zero. Rather, it suggests that the negative contributions are more prevalent and tend to, on average, be smaller in magnitude. This implies that the distribution over *D*_0_ values is skewed — and the CP metrics may serve as useful metrics for understanding that skew.

**Figure 3:**
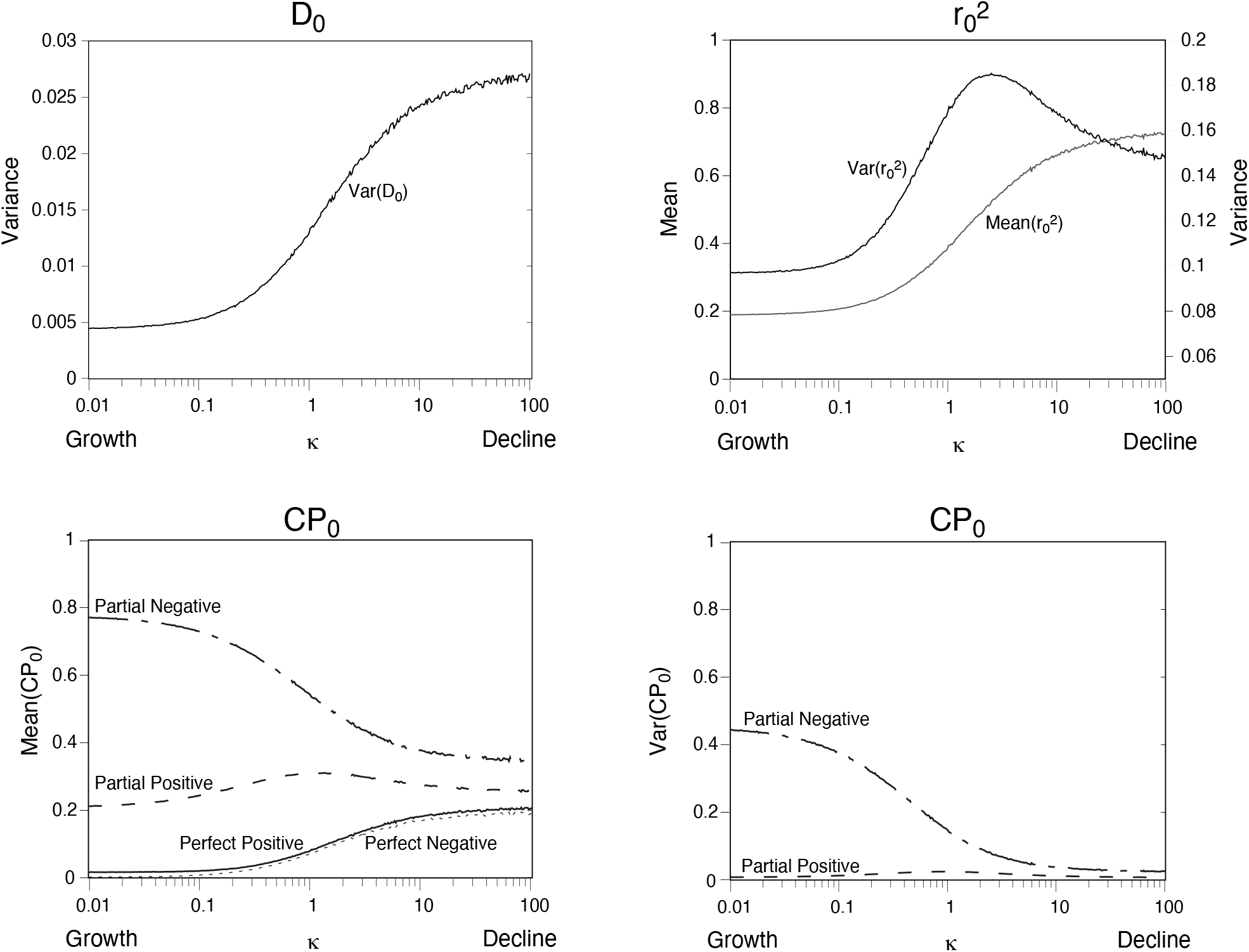
How LD metrics (*D*_0_, 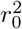 and CP_0_) vary with demography (*κ* on x-axis), even without recombination.

#### 3.1.3 Demography

##### Distribution of *D*_0_

To get a sense of how *D*_0_ is changed by the demography, we generated Monte-Carlo distributions of the expected value of *D*_0_ given the genealogy, across trees, without recombination. We assumed that ancestral states are known (e.g., using an outgroup and no orientation errors), so that allele frequencies at both loci correspond to true derived-allele frequencies. For each demographic scenario, we simulated trees and computed all possible covariances of allele frequencies arising from two independent mutations placed on the same genealogy, thereby obtaining the full distribution of *D*_0_ over the entire tree space. As expected from our analytical results, we observe that *D*_0_ has a mean of zero regardless of the demographic scenario.

In Table 1 we report the variance of *D*_0_ under three different demographic scenarios: increasing population (*κ* = 0.5), constant population (*κ* = 1) and decreasing population (*κ* = 2); considering three different number of leaves per tree: *n* ∈ [10; 50; 100]. We find that the variance of *D*_0_ decreases as either the number of sampled leaves increases or the population size decreases. The variances obtained from simulations are consistent with those computed using Equation 2.

**Table 1:** Variance of *D*_0_ depending on the number of leaves in the tree (n) and the strength of change *κ* with a time of *τ* = 0.5, estimated on 1,000 trees.

| $n$ | $\kappa = 0.5$ | $\kappa = 1$ | $\kappa = 2$ |
| --- | --- | --- | --- |
| 10 | 0.0088 | 0.0117 | 0.0150 |
| 50 | 0.0029 | 0.0049 | 0.0078 |
| 100 | 0.0021 | 0.0038 | 0.0066 |

Figure 2 shows the full distributions of *D*_0_ for the case *n* = 100. The top panels correspond to these first simulations, where ancestral states are known and derived allele frequencies can be used directly. We also computed 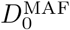, the covariance of the two minor allele frequencies. Its distribution is shown in the bottom panels of Figure 2.

As stated above, *D*_0_ has mean zero under all demographic scenarios. In contrast, the MAF-based statistic 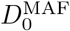 exhibits a slightly positive mean, consistent with previous observations (Good, 2022).

##### Influence of demography parameters on LD measures

The variance of *D*_0_ depends on the demography, with large variances for declining populations and small variances for growing populations, pointing to its potential usefulness in assessing demography. We therefore explored more extensively the impact of the demography on the patterns of linkage by varying the parameter *κ* ∈ [0.01, 100]. Because there is no recombination, |*D*′|_0_ = 1 for possible values of *κ*. We thus only report the variance of *D*_0_, the mean and variance of 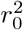 and CP_0_.

Results (Figure 3) confirm that the variance of *D*_0_ and the mean of 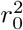 exhibit highly similar behaviour across demographic scenarios. Because E[*D*_0_] = 0, we have 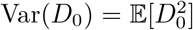, which is precisely the numer-ator of 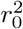. Thus, apart from the normalization in the denominator, both statistics respond in parallel to demographic changes, and we indeed find that both Var(*D*_0_) and 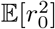 are larger in declining populations than in expanding ones.

This pattern is a direct consequence of how demographic variation reshapes coalescent trees. Expanding populations produce genealogies with short internal branches and comparatively long external branches, whereas declining populations generate trees with extremely long root branches and clearly defined subtrees. These distortions affect where mutations fall on the tree and therefore whether they generate positive or negative configurations of *D*_0_.

In expanding populations, long external branches increase the chance that mutations arise in parts of the genealogy where numerous subtrees coexist, which favours configurations leading to values of *D*_0_ close to zero and therefore a small variance. In contrast, declining populations have long internal branches descending from the root that split the sample into two deep subtrees. These well-separated subtrees mechanically increase the proportion of perfect positive and perfect negative configurations (in terms of concordant or discordant mutation pairs), leading to a higher proportion of these CP_0_ outcomes in declining populations than in expanding ones. Mutations occurring on the same branch tend to produce large positive *D*_0_, while mutations falling on different branches generate strongly negative values. The mixture of these outcomes leads to a widespread and therefore large variance of *D*_0_.

These differences in tree shape are typically captured by the site-frequency spectrum (SFS) (Fu, 1995), which is widely used for demographic inference (Gutenkunst *et al*., 2009; Excoffier *et al*., 2013; Liu and Fu, 2020). Here, instead of allele counts, we observe the covariance between pairs of mutations. Expanding populations, dominated by low-frequency variants, necessarily concentrate *D*_0_ values near zero, whereas declining populations—with more intermediate- and high-frequency variants—produce a broader range of *D*_0_ values.

Results obtained using 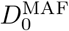 displayed the same qualitative behaviour (see Supplementary Figures 12, 13, 14).

##### Comparison to two-locus moment expectations

Linkage disequilibrium has traditionally been studied using analytical, moment-based approaches, including two-locus statistics derived under a variety of scenarios (Lewontin, 1964; Hill and Robertson, 1968; Ohta and Kimura, 1969a; Sved, 1971; Hill and Weir, 1988). To benchmark our simulation framework, we focus on 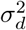 and compare the results from our coalescent simulations to analytical predictions obtained via the two-locus moment recursion method (Hill and Weir, 1988).

The comparison is conducted under the same demographic scenario: a sudden population contraction parameterized by the contraction strength *κ* and the time since contraction *τ*. Analytical expectations for 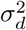 are obtained by solving a system of recursion equations for the relevant two-locus moments, including ⟨*p*_*i*_*q*_*i*_⟩, ⟨*p*_*i*_*q*_*i*_*p*_*j*_*q*_*j*_⟩, ⟨*p*_*i*_*p*_*j*_*D*_*ij*_⟩, and 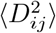, and expressing the dynamics in coalescent time units (Supplementary Material 7). The equilibrium values for a constant population size recover previously reported results (Ohta and Kimura, 1971; Hill and Weir, 1988), and the system can be numerically solved under varying *κ* and *τ* to capture the effects of demographic change. Multiple values of *τ* are explored in Supplementary Figure 11.

In our simulations, LD is computed for trees weighted by their total branch length. Comparing simulation results to analytical predictions shows strong concordance, after accounting for the factor-of-two difference in *τ* between haploid (simulation) and diploid (analytical) frameworks (Fig. 4). These results indicate that coalescent simulations weighted by tree length provide an accurate representation of two-locus moment expectations under both constant and changing population sizes.

**Figure 4:**
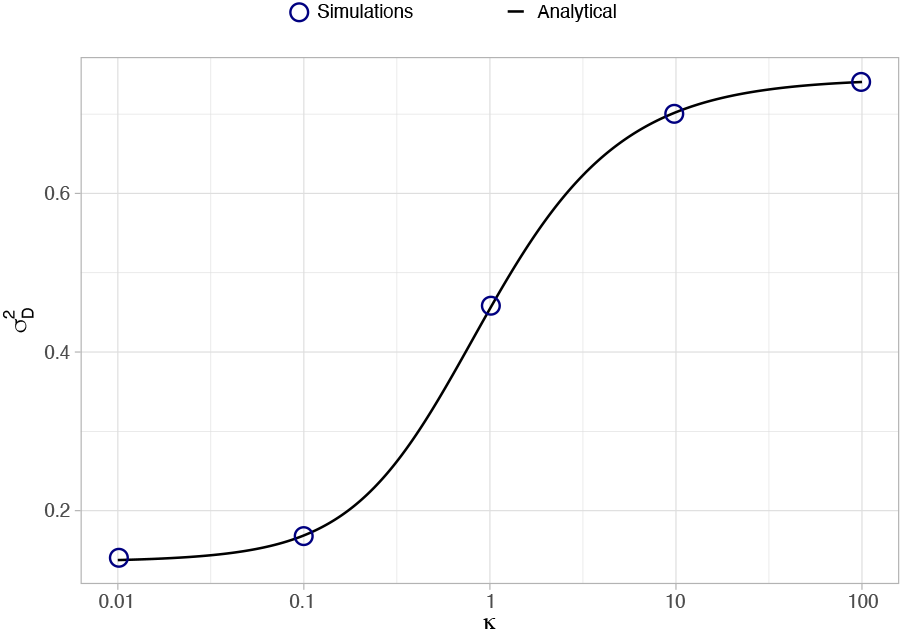
Estimates of 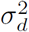 for varying *κ* ∈ [0.01, 0.1, 1, 10, 100]. Coalescent simulations are performed with *τ* = 0.5 and *n* = 500, averaging results over 5000 simulated trees. For the two-locus moments estimate, we set *τ* = 1 to take into account the ploidy, as explained in the main text.

Importantly, despite relying on two fundamentally different methodologies: one based on solving recursive expectations for two-locus moments, the other on explicit genealogical simulations; they are both designed to capture the same underlying process, and accordingly yield consistent qualitative and quantitative trends across demographic scenarios. This agreement shows that the two perspectives are coherent within a common coalescent framework and supports the interpretation that the observed patterns reflect the effects of demography on LD.

### 3.2 With one recombination event

#### 3.2.1 Classification of Recombination events

To study the effect of a *single* recombination event, we first need to define the four different types of recombination events, depending on their effect on the tree topology. Indeed, following the sequential approach to study recombination (Wiuf and Hein, 1999), the coalescent tree of the locus on one side of the recombination event (here locus 1) may be different from the tree of the locus on the other side of the recombination event (here locus 2).

##### Definitions

Considering the effect of a single recombination event on a coalescent tree, we have defined 4 recombination types :

**Hidden :** No tree change. The recombination event and the coalescent event happened on the same lineage (Figure 5a).

**Figure 5:**
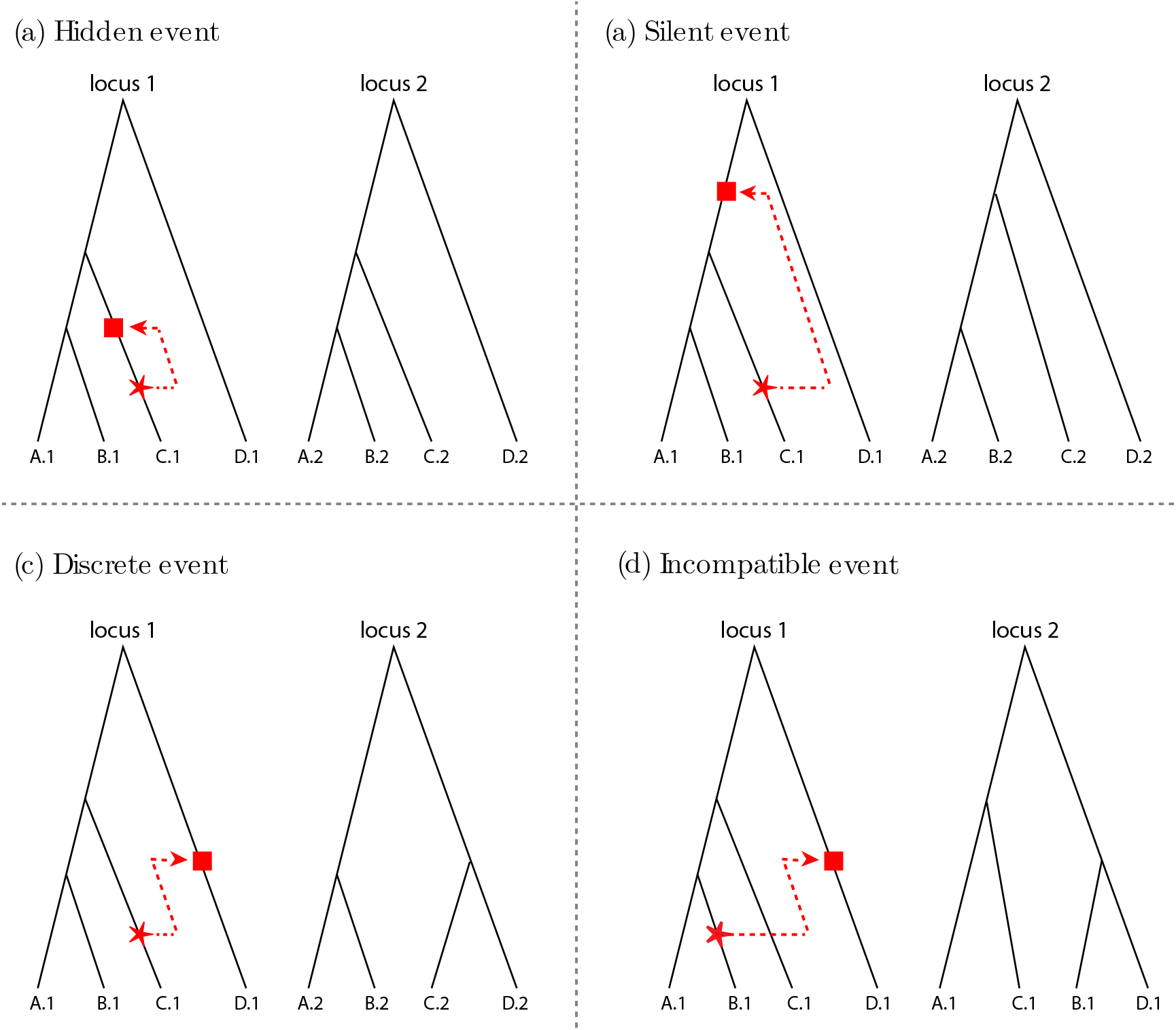
A sketch of the trees at the two loci on both sides of the recombination event (locus 1 on the left and locus 2 on the right). The red arrow indicates the recombination event affecting the genealogy at locus 1, while the red square marks the point of re-branching in the tree resulting in the locus 2 tree. The direction of the arrow illustrates how the recombination event may or may not alter the tree topology when moving from locus 1 to locus 2. Recombination events can be sorted depending on their effect on the topology change (from locus 1 to locus 2). We consider four types of recombination events: a) hidden events result in no change in the tree (they cannot be detected), b) silent events only affect the branch length (so relative abundance of variants could change), c) discrete events change only the rooted tree (without an outgroup, it would be a silent event), while d) incompatible events change the unrooted tree (and thus are the easiest to detect).

**Silent :** Only change in the branch lengths, not in the tree topology (Figure 5b). The recombination event happened with an ancestor of the recombining lineage.

**Discrete :** Change in topology and branch lengths. However, the un-rooted topology does not change. This type of recombination is not detectable using the Four Gamete Test (Hudson and Kaplan, 1985), but can be detected if the ancestral alleles are known. This type of recombination happened when the recombining lineage coalesces with a lineage with the same ancestor (Figure 5c).

**Incompatible :** Important topology change, detectable using the Four Gamete Test. This type of recombination happened when the recombining lineage coalesces with a lineage with a different ancestor (Figure 5d).

These recombination types are defined as in Ferretti *et al*. (2013) with a sub-division of Non-silent recombination into Discrete and Incompatible. Hidden recombination correspond to ‘type-1’ in Deng *et al*. (2021), Silent to ‘type-2’ and ‘type-3’, and Discrete and Incompatible recombinations combined to ‘type-4’.

##### Proportions

The proportions of the different types of recombination events will highly depend on the coalescent tree.

First, the number of leaves, which are a direct output of the sampling effort, will change the different proportions. For example, for trees with less than 4 leaves, there are no possible incompatible events, whereas they are likely more abundant in dense trees with many branches. As illustrated by Figure 6, the proportions of incompatible recombination events indeed increases with the number of leaves of the tree, while the silent ones becomes rarer. In a coalescent tree with many leaves, as there are many branches (whose average lengths tend to be small), there are less opportunities for silent events to occur.

**Figure 6:**
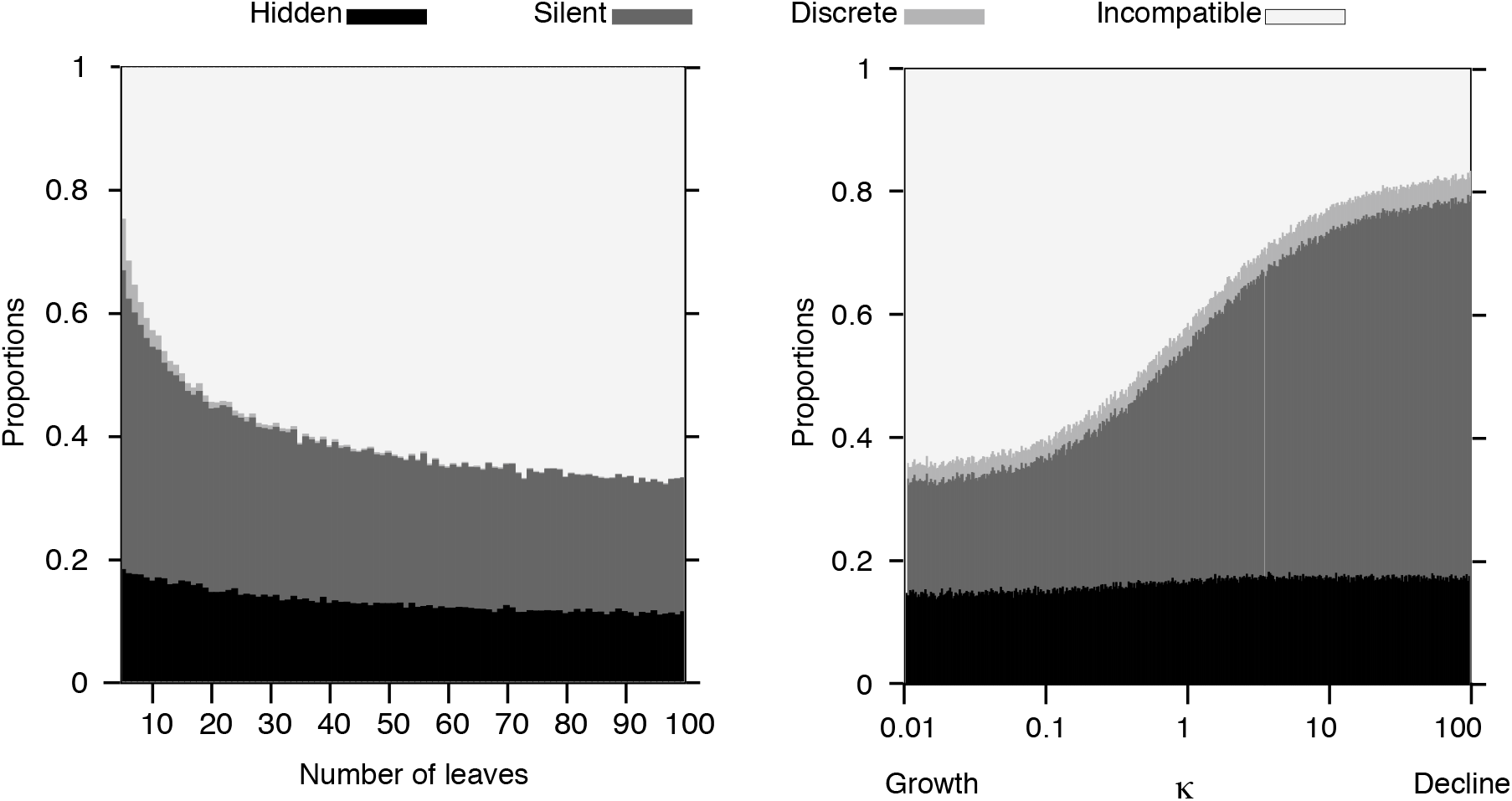
How the proportions of the four types of recombination events vary with number of leaves (left panel, *κ* = 1) or demographic variations (right panel, leaves *n* = 10).

Second, the different events will also depend on the exact tree topology and branch lengths. As they both vary a lot with the population demography, we expected that the four types of events will have different proportions according to the demography. For example, in a declining population, the coalescent tree has long root branches where recombination events can only be silent ones. On the contrary these root branches are short in growing populations, leaving fewer opportunities to silent events. As illustrated by Figure 6, incompatible and silent events are strongly affected by the overall demography of the population.

Interestingly, the proportions of Hidden recombination events are not really impacted by either the density of the tree or by the demography of the population (Figure 6).

#### 3.2.2 Influence of demography parameters and type of recombination on LD measures

*D*_1_ **and** 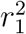: The variance of *D*_1_ and the mean of 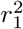 vary similarly. The general trend is the same as *D*_0_ and 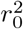: low for the increasing population and high for the declining population. The four types of recombination can be divided in two groups: affecting topology (Discrete and Incompatible) and no effect on topology (Silent and Hidden). When the topology is affected, the two trees are less correlated and the magnitude of LD is lower than in the absence of recombination: *D*_1_ *< D*_0_ and 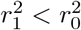 (Figure 3 and Figure 7). On the other end, when the topology is not affected, the two trees are highly correlated and the LD measures are similar than without recombination: *D*_1_ ≈ *D*_0_ and 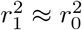 (Figure 3 and Figure 7).

**Figure 7:**
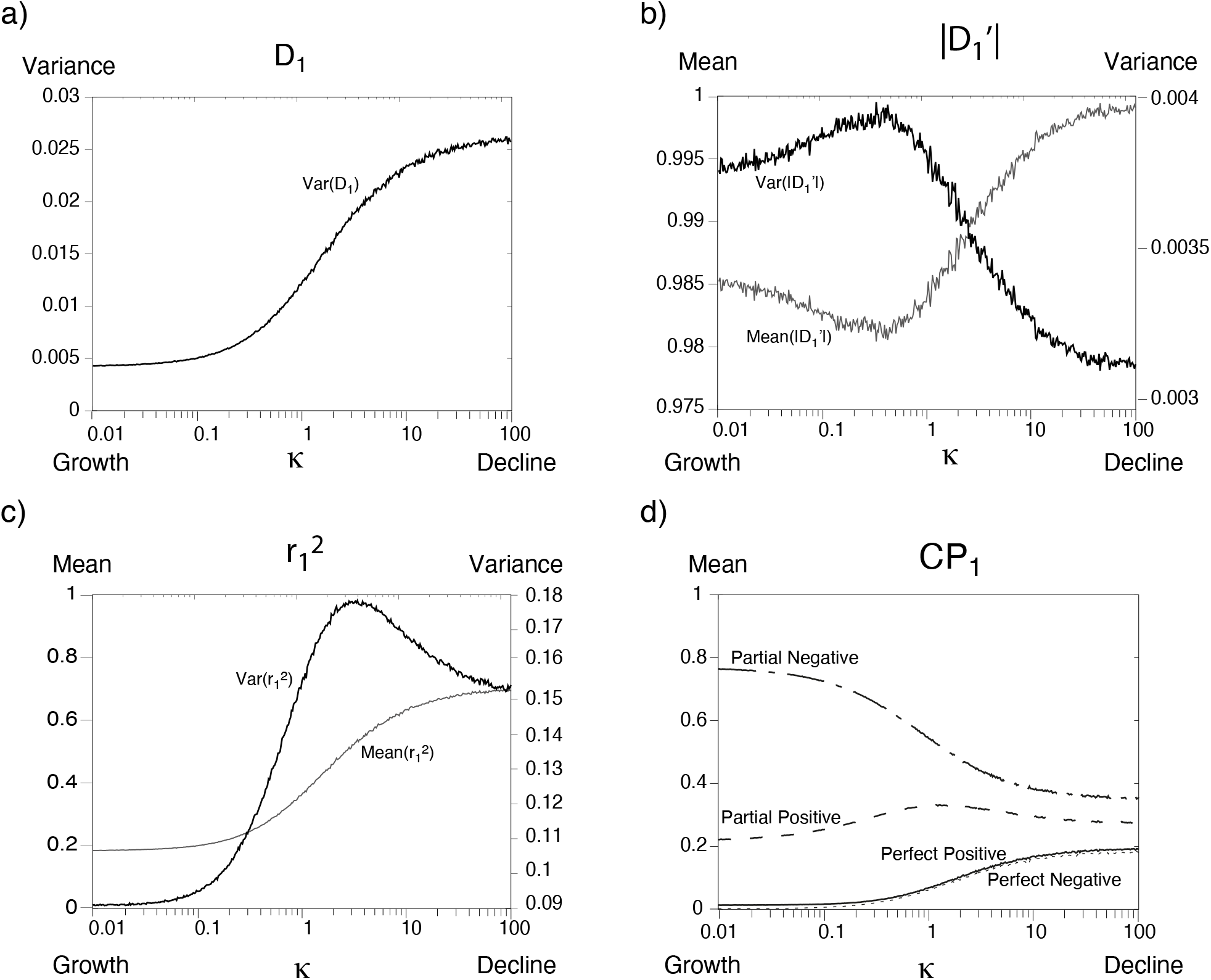
LD measures *D*_1_ (all recombination types) (a), 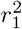 (b), |*D*′|_1_ (c) and CP_1_ (d) between alleles on two adjacent trees, in function of the strength of contraction (x-axis).

The difference in *Var*(*D*_1_) between Discrete and Incompatible recombination for declining populations can be explained by the branch length proportion affected by the topology change. Discrete recombination happened in more ancestral branches than Incompatible recombination and these branches are really long for a declining population. Less long-length branches are correlated after a Discrete recombination event than after an Incompatible recombination event and *Var*(*D*_1_) is lower for Discrete recombination. 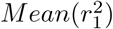 does not vary between Discrete and Incompatible recombination for a declining population, but does for an increasing population. The exact opposite phenomena happens in an increasing population, as ancestral branches are shorter. The most recent branches, where many Incompatible recombinations have the chance to happen, are more important in the total branch length and thus, in the correlation between trees.

|*D*′|_1_: The value of |*D*′|_1_ is highly influenced by the recombination type, as only the Incompatible recombination events generate |*D*′|_1_≠ 1. However, if only one recombination event happened, no more than two branches will be incompatibles. The mean value of |*D*′| is impacted by the number of incompatibilities generated and the proportion of the incompatible branches on the total tree length. For large declines in population, the ancestral branches represent the large majority of the total tree length and cannot generate incompatibilities. In that case, an Incompatible recombination event affects a low proportion of the total tree length, and the mean |*D*′|_1_ is not at its maximum (Figure 7). It will be at its maximum for a well-balanced tree, around *κ* = 1 (Figure 7).

**CP**_1_: We also examine the 2-site configuration probabilities between trees separated by each recombination type. Hidden and silent recombination events tended to have slightly fewer partial negative configurations, and slightly more frequent perfect configurations, as shown in Figure 8, the opposite pattern was seen for discrete and incompatible recombination events, with the mean partial negative configuration being larger and the perfect negative fractions decreasing. The same pattern was observed across demographic scenarios.

**Figure 8:**
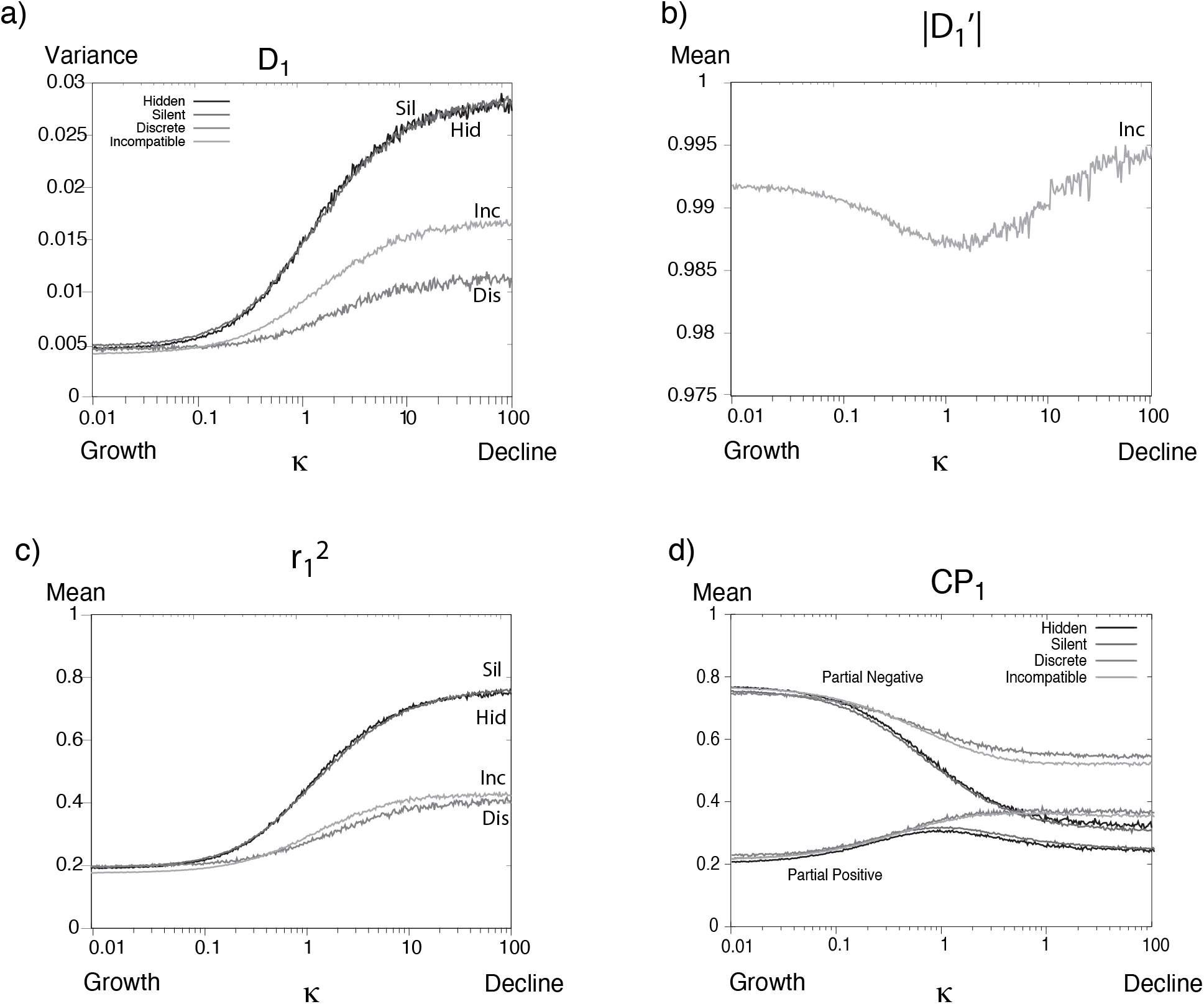
LD measures *D*_1_ (a), 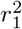, |*D*′|_1_ (c) and CP_1_ (d) between alleles on two adjacent trees, in function of the strength of contraction (x-axis). Different line types indicate the different type of recombination. Only Partial Negative and Partial Positive CPs categories are displayed, as Perfect Negative and Perfect Positive values are less variable.

### 3.3 With an infinite number of recombination events - Two independent trees

Results for an infinite number of recombination events are reported on Figure 9.

**Figure 9:**
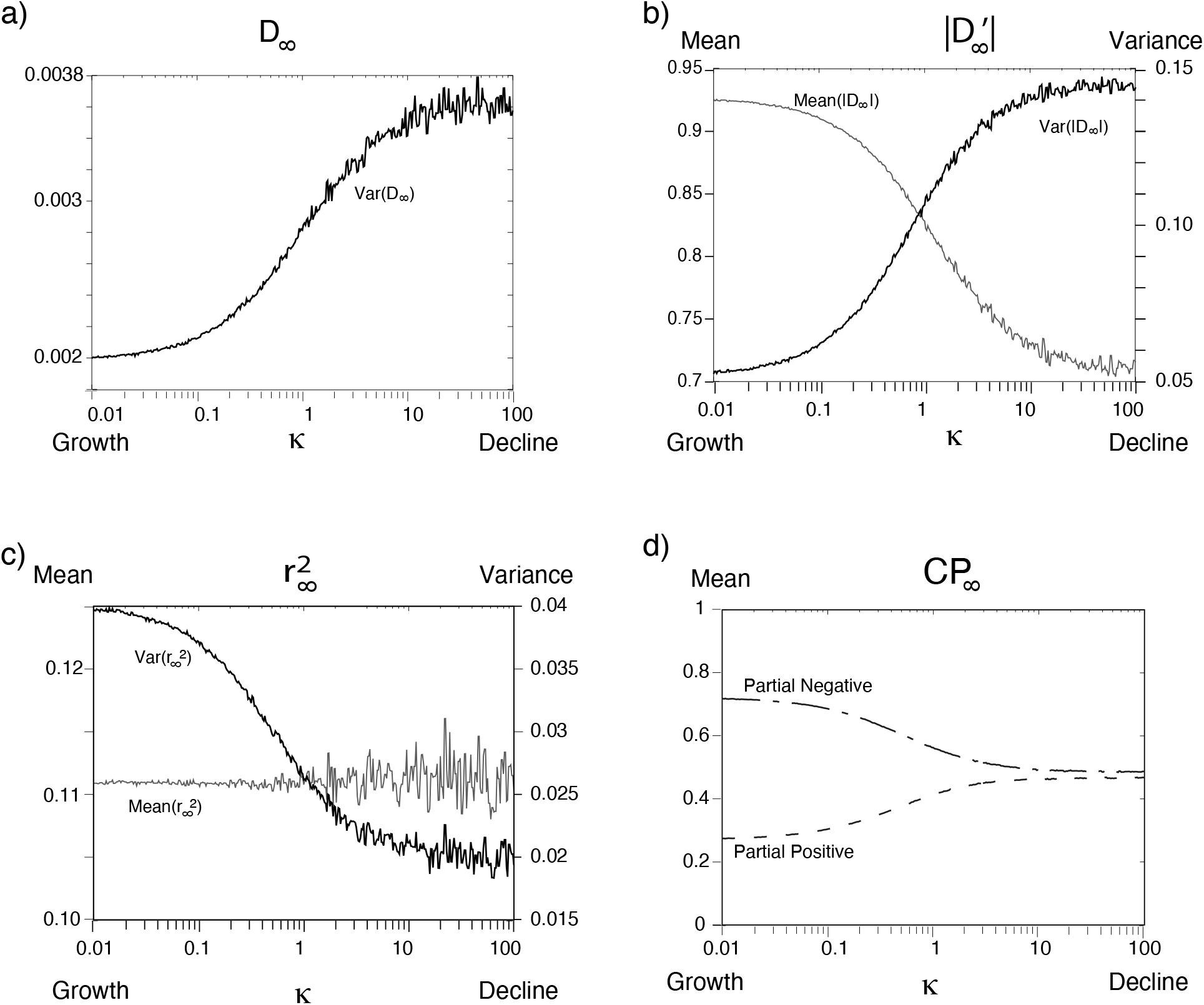
LD measures *D*_∞_ (a), 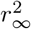, |*D*′ | (c) and CP_∞_ (d) between alleles on two independent trees, in function of *κ* the strength of contraction (x-axis). Only Partial Negative and Partial Positive CPs categories are displayed, as Perfect Negative and Perfect Positive values are nearly negligible.

**Figure 10:**
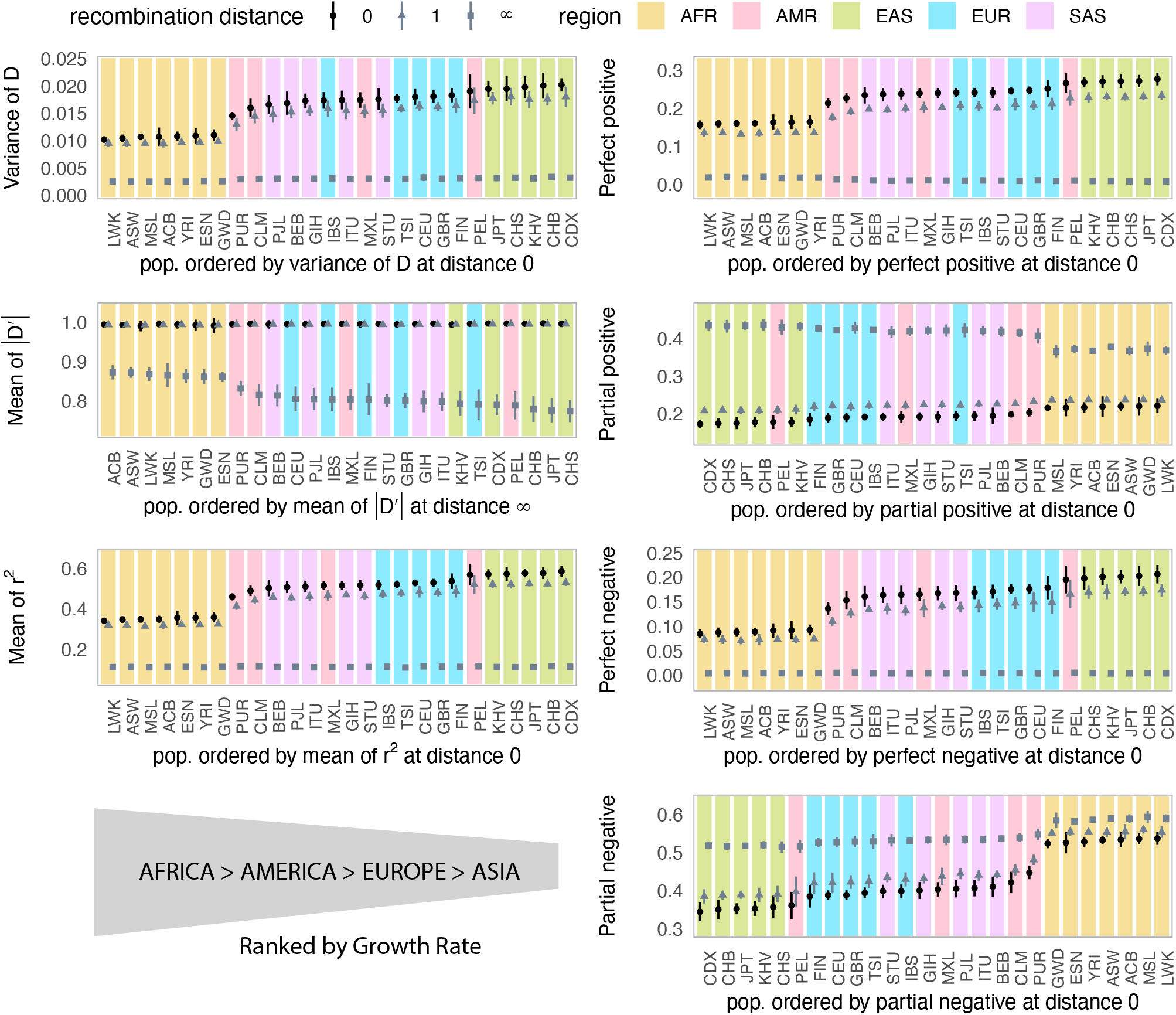
LD estimates across recombination scales using 1KG3 tree sequences. LD measures were calculated from phased genotype data at three different recombination scales: 0, within the same tree (circles); 1,between adjacent trees (triangles), and ∞, approximated by using tree pairs drawn from different chromosomes (squares). For each 1KG3 group, we report mean values and 95 % confidence intervals across ten independent sample sets of ten haploid genomes. Colors indicate continental regions: Sub-Saharan Africa (AFR, orange), Americas (AMR, red), East Asia (EAS, green), Europe (EUR, blue), and South Asia (SAS, pink). Populations are ordered by their mean LD values to highlight broad-scale variation patterns. Each tree contributed equally to the overall distribution, with averages weighted by the number of mutation pairs per tree.

**Figure 11:**
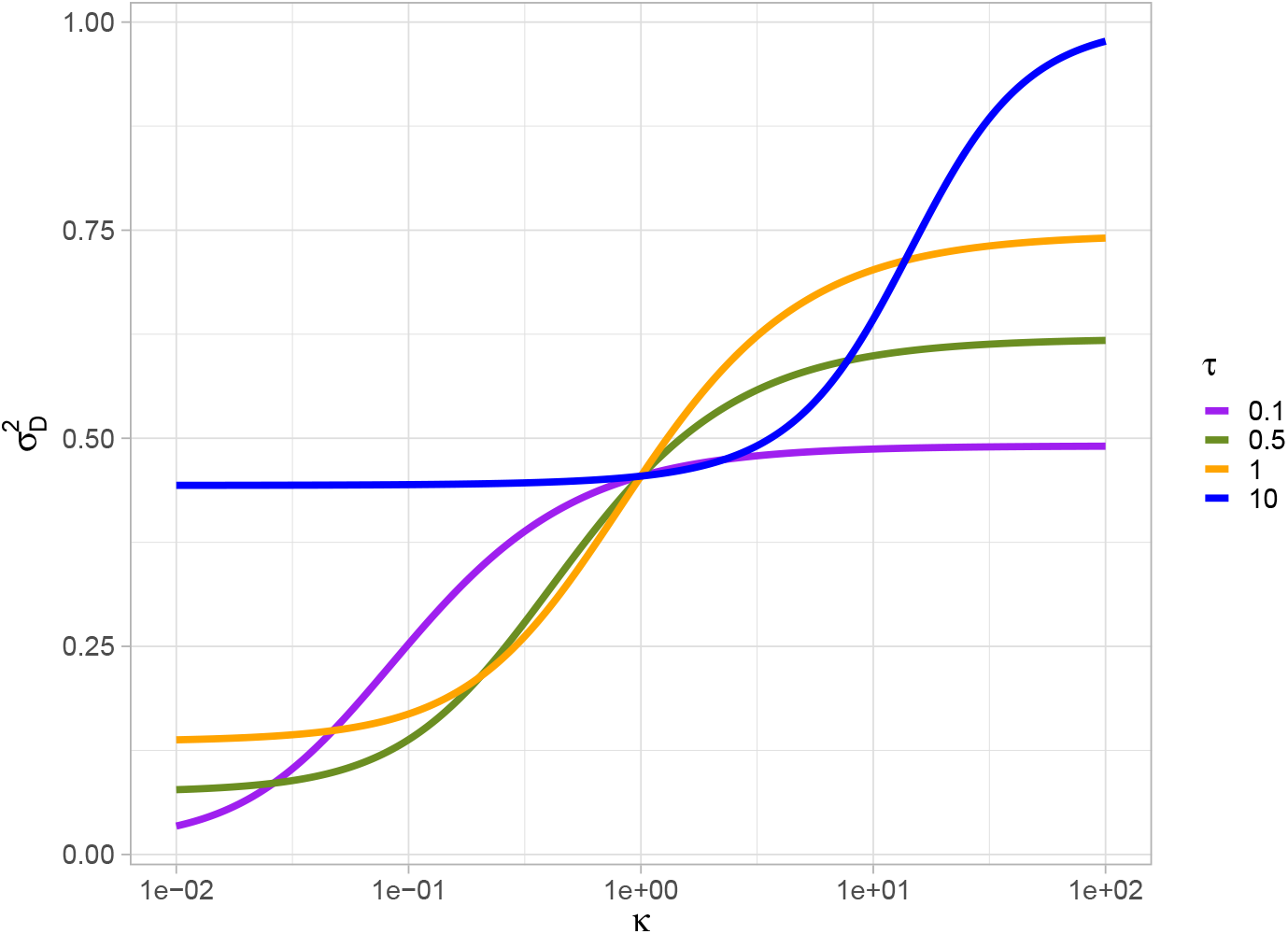
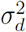 with varying *κ* ∈ [0.01 : 100] (x-axis) for *τ* ∈ [0.1, 0.5, 1, 10] (different colors). For a constant population size scenario (*κ* = 1), 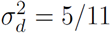.

**Figure 12:**
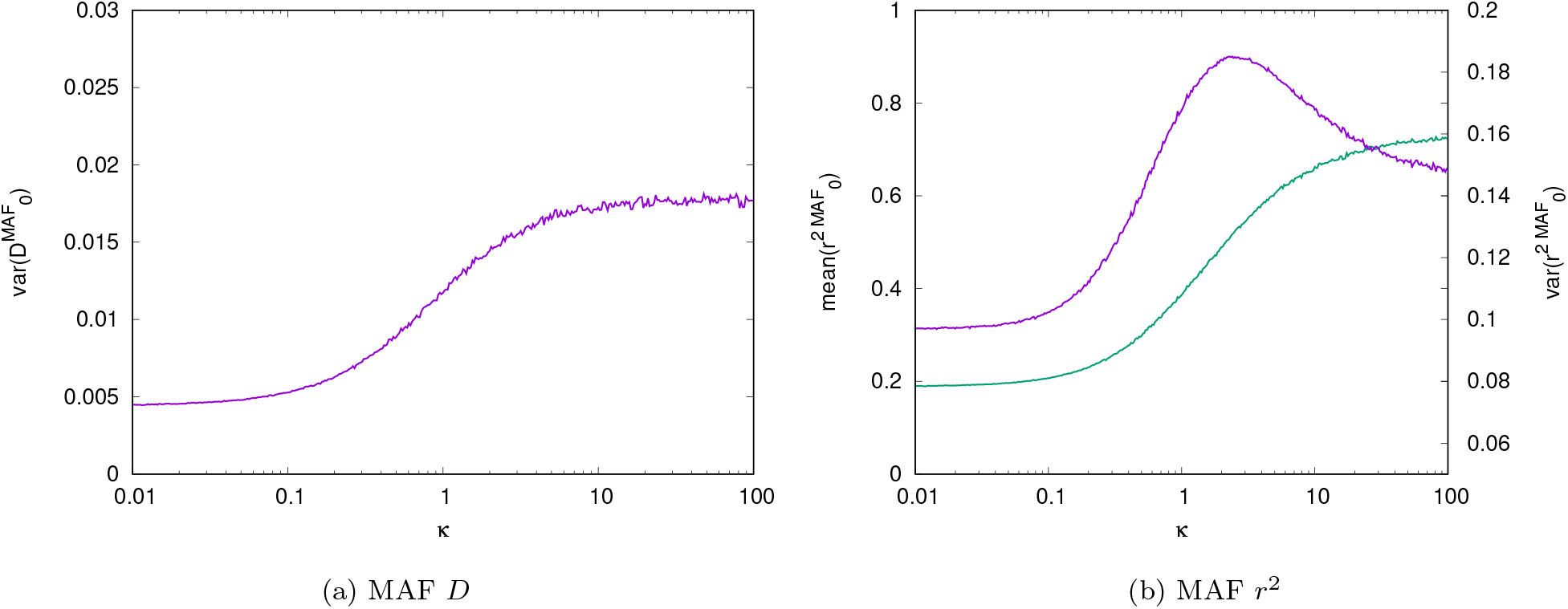
MAF LD measures *D* (12a) and *r*^2^ (12b) between alleles on the same tree, in function of the strength of contraction (x-axis). (12a) *Mean*(*D*) in green and *Var*(*D*) in purple, (12b) *Mean*(*r*^2^) in green and *Var*(*r*^2^) in purple.

**Figure 13:**
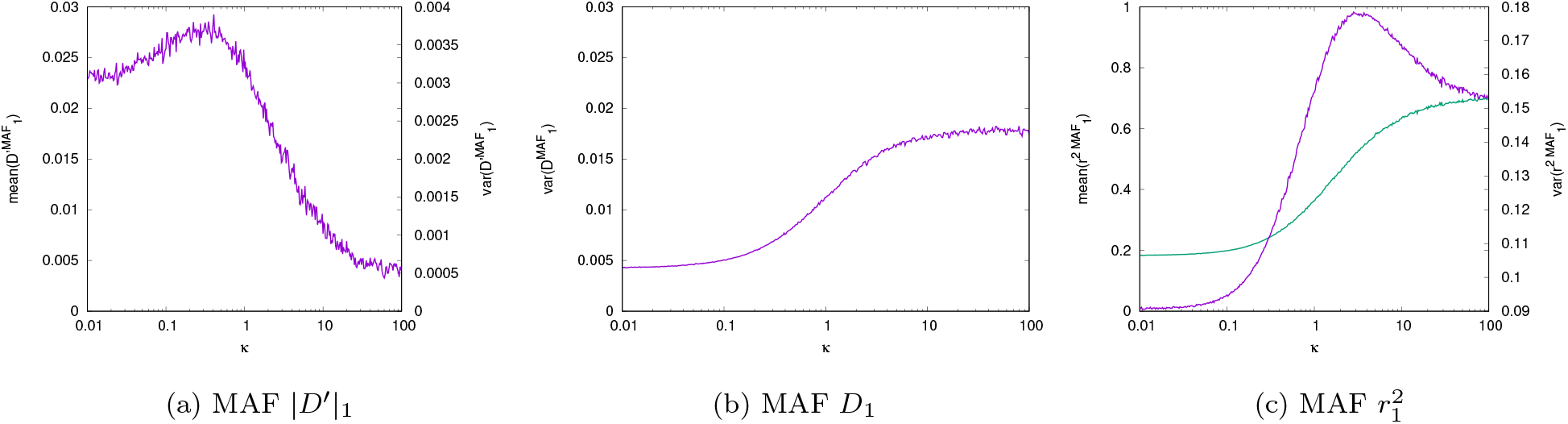
MAF LD measures |*D*′| (Figure 13a), *D* (13b) and *r*^2^ (13c) between alleles on two different trees, in function of the strength of contraction (x-axis). (13b) *Var*(*D*) in purple, (13c), *Mean*(|*D*′|) in green and *Var*(|*D*′|) *Mean*(*r*^2^) in green and *Var*(*r*^2^) in purple.

**Figure 14:**
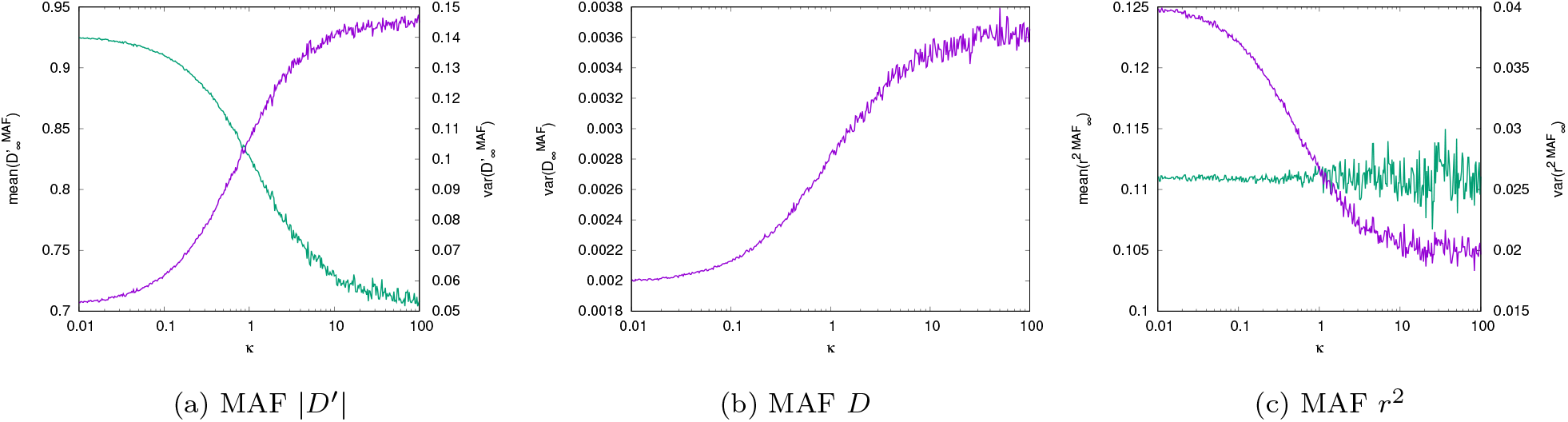
MAF LD measures |*D*′| (Figure 14a), *D* (14b) and *r*^2^ (14c) between alleles on two different trees, in function of the strength of contraction (x-axis). (14b) *Var*(*D*) in purple, (14c), *Mean*(|*D*′|) in green and *Var*(|*D*′|) *Mean*(*r*^2^) in green and *Var*(*r*^2^) in purple.

*D*_*∞*_: *Var*(*D*_∞_) in different demographic scenarios has the same behavior as *Var*(*D*_0_): low for increasing-population-size trees and high for decreasing-population-size trees. However, the range of value is different, *Var*(*D*_∞_) *< V ar*(*D*_0_). The highest value of *Var*(*D*_∞_) is lower than the lowest value of *Var*(*D*_0_). Recombination events break linkage between sites and thus, decrease *Var*(*D*). *Var*(*D*) decreases with the number of recombination events between alleles (not shown).

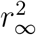 : In the case of two independent trees, 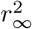 does not vary with demography. The normalization by allele frequency reduces the effect of demography for unlinked trees.

|*D*′|_*∞*_: The *Mean*(|*D*′|_∞_) varies with the demography. *Mean*(|*D*′|_∞_) is low in declining population trees and high in increasing population trees. Trees with long ancestral branches (e.g. decreasing population tree) will, proportionally, be less incompatible than trees with short ancestral branches (e.g. increasing population tree). Even independent trees will not have striking incompatibilities if simulated under a strong decline in population.

**CP**_*∞*_: Because the two trees are independent, the number of perfect positive and perfect negative configurations is negligible. Focusing on partial positive and partial negative configurations, we observe similar trends with respect to demography: the proportion of partial negative configurations decreases as the population declines, while the proportion of partial positive configurations increases under the same conditions.

### 3.4 LD-measures and number of recombination events

Comparison of all three statistics for 0, 1 or an infinite number of recombination events are shown in Figures 16 & 17.

**Figure 15:**
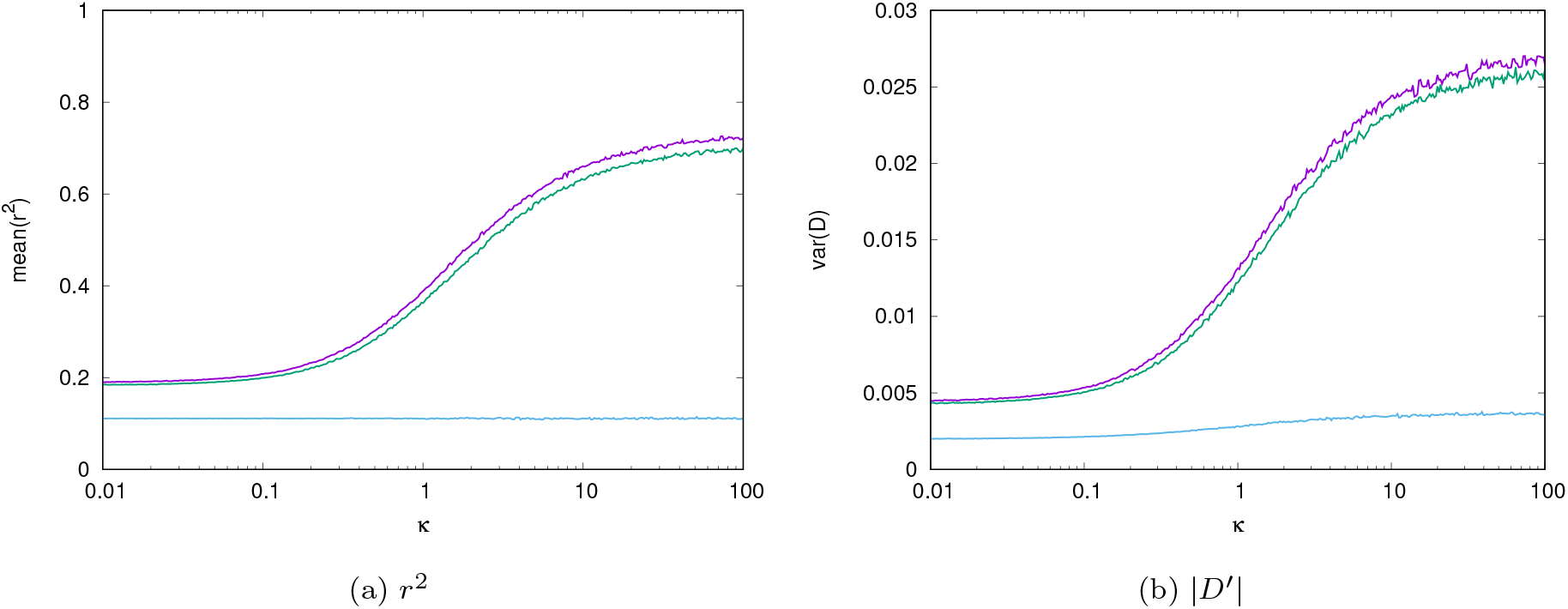
LD measures *r*^2^ (Mean) and |*D*′| (Variance) in function of the strength of contraction (x-axis), for the three different recombination scenarios: no-recombination in purple, one recombination events between the alleles used to compute the LD measure in green and the alleles on two independent trees in blue.

**Figure 16:**
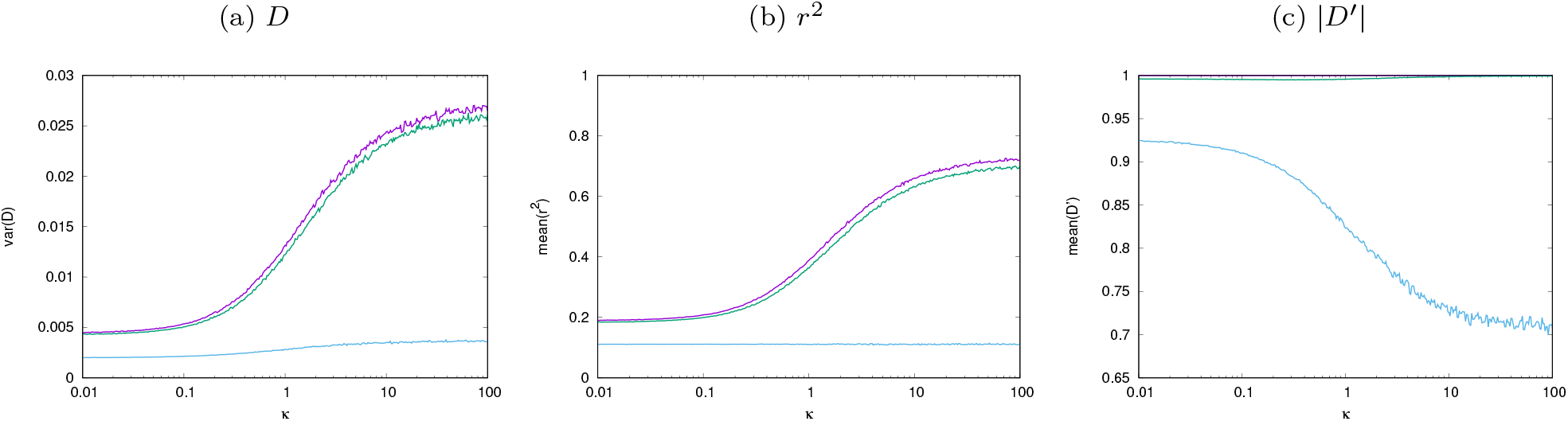
LD measures *D* (Variance), *r*^2^ (Mean) and |*D*′| (Mean) in function of the strength of contraction (x-axis), for the three different recombination scenarios: no-recombination in purple, one recombination events between the alleles used to compute the LD measures in green and the alleles used to compute the LD measures on two independent trees in blue.

**Figure 17:**
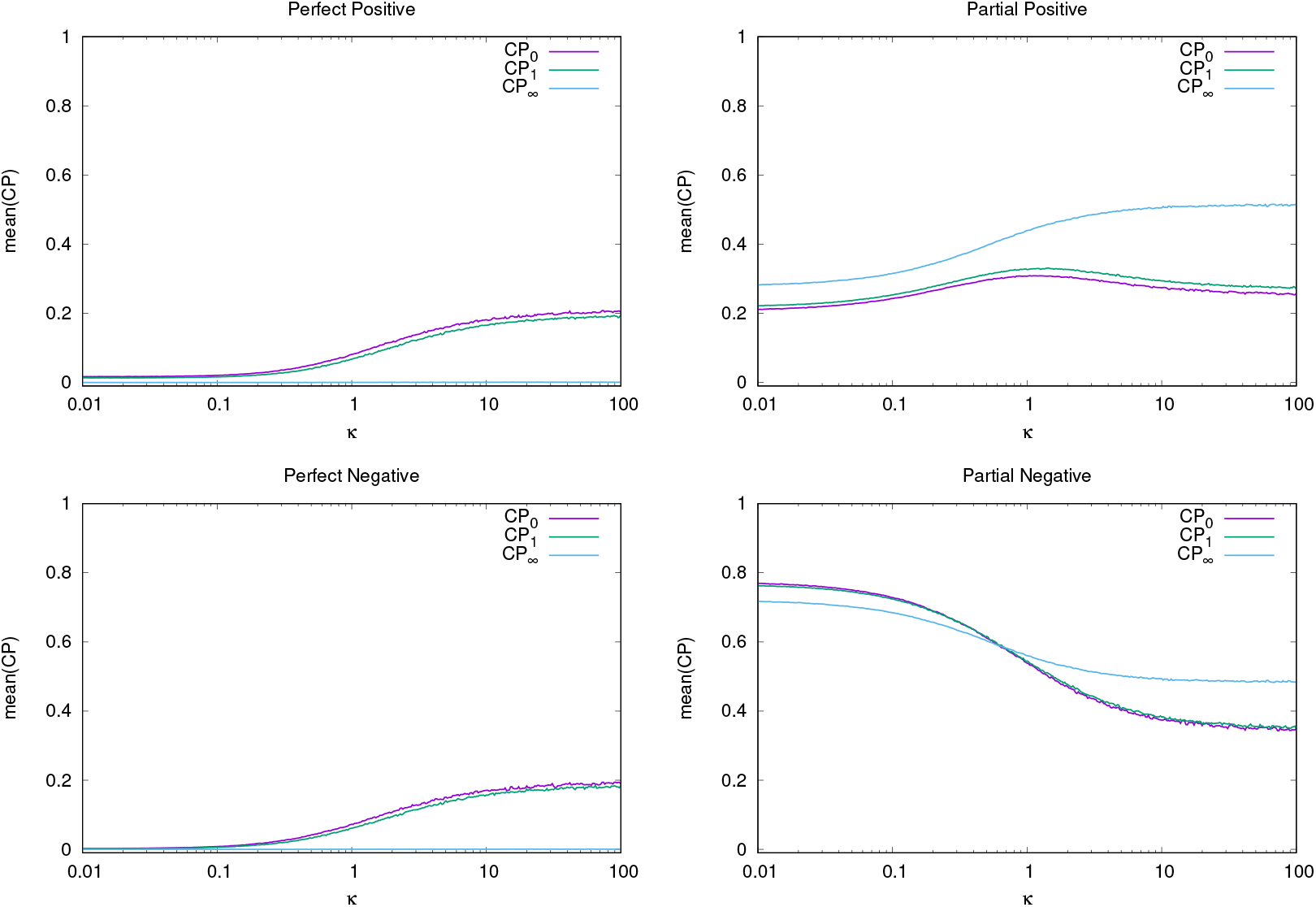
2-site configuration probabilities in function of *κ* the strength of contraction (x-axis), for the three different recombination scenarios: no-recombination, one recombination events between the alleles used to compute the LD measures and two independent trees.

*D*: *Var*(*D*) has the same behavior depending on the demographic scenario: lower for increasing population trees and higher for decreasing population trees. However, the range of value change drastically with the recombination scenario considered. We observe *Var*(*D*_∞_) *< V ar*(*D*_[0−1]_), varying from two times smaller (for *κ* = 0.01, *Var*(*D*_0_) = 0.0047 and *Var*(*D*_∞_) = 0.002) for an increasing population to nearly ten times smaller (for *κ* = 125, *Var*(*D*_0_) = 0.026 and *Var*(*D*_∞_) = 0.003) for a decreasing population. On the other hand *Var*(*D*_0_) and *Var*(*D*_1_) are similar, even for extreme values (for *κ* = 0.01, *Var*(*D*_1_) = 0.0045 and for *κ* = 125, *Var*(*D*_1_) = 0.025) (Figure 16a).

*r*^2^: As said previously, 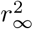 does not vary with demography. This is strikingly different from the the case with zero or one recombination in which *r*^2^ increases with decreasing population sizes. As we can see in Figure 16b, the mean of 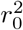 and 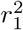 are similar. However, we want to note that there is a shift in the variance when considering two alleles on the same tree or on two trees separated by one recombination event 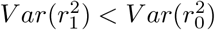 (Figure 15a).

|*D*′|: For both the mean and the variance the only |*D*′| varying significantly with the demography is |*D*′|_∞_ (Figure 16c,15b). The trend of |*D*′| is the opposite of *D* and *r*^2^, as it’s decreasing with decreasing population size.

**CPs:** 2-site configuration probabilities vary strongly across demographic scenarios. Compared to populations of constant size, growing populations tend to have more frequent negative configurations, compared to positive, and less frequent perfect configurations overall (Figure 17). Shrinking populations show an opposite pattern, with similar frequencies of partial negative and partial positive configurations and overall more frequent perfect configurations. The trend is virtually identical in trees separated by one recombination event. In trees separated by infinitely many recombination events, the frequencies of the perfect configurations are tiny, while those of the partial configurations are closer to each other.

### 3.5 Application

Tree sequences provide a natural framework for measuring statistics as a function of the number of detectable recombination events. The genomic span represented by a single tree captures the scale of genealogical linkage disequilibrium. Neighboring trees are separated by one recombination event, whereas trees drawn from different chromosomes can be thought of as separated by an infinite number of recombination events. When applying these ideas to empirical data, however, we found that tree-based LD estimates are subject to substantial biases, particularly when contrasted with estimates obtained from true genealogies (Figure 19). To reduce the impact of such inference-related artifacts, we instead rely on genotype-based LD estimates, computed directly from phased variants but restricted to the genomic spans defined by the inferred trees. This strategy preserves the genealogical resolution provided by the tree-sequence framework while avoiding the distortions inherent in the tree-based LD calculations.

All the LD statistics measured generally fall into the range observed in our simulations and analytical predictions. Sub-Saharan African (AFR) populations tended to have lower *r*^2^ and *Var*(*D*), but higher |*D*′|, consistent with a more growth-like demographic history. In contrast, East Asian (EAS) populations evidence the opposite pattern, suggesting demographic shrinkage, with American (AMR), European (EUR) and South Asian (SAS) populations falling in between. This ordering remained consistent across all measured statistics. A similar pattern emerged for the 2-site configuration probabilities, where AFR populations had lower perfect positive and perfect negative configuration probabilities, but higher partial negative configuration probability. The ordering of partial positive configuration probabilities at distance 0 was less consistent, reflecting the non-monotonic relationship of this statistic with *κ*, which limits its interpretive value for demographic inference.

Comparing the estimates to our simulated simple ‘single population size change’ scenario as in our simulations, most statistics suggest population growth (*κ <* 1) in Sub-Saharan Africans and population decline (*κ >* 1) in non-African populations, consistent with the Out-of-Africa model of human history. Although this simplified model does not capture the full complexity of demographic processes, it provides a useful benchmark to identify which statistics are informative and which deviate from model predictions.

For example, under the infinite recombination scenario tested here (*κ* ∈ [0.01, 100], *τ* = 0.5 and *n* = 10) Perfect Positive and Perfect Negative configurations are expected to occur only at negligible frequencies. Instead, empirical frequencies are markedly higher than even the maximum simulated values: Perfect Positive ≈ 0.01 vs. *<* 0.003; Perfect Negative ≈ 0.004 vs. *<* 0.00125. (These simulated frequencies were too low to be visible in Figure 9.) Likewise, in the single recombination event model, very high values of |*D*′|_1_ are expected. However, because invisible or silent recombination events are undetectable, the empirical |*D*′|_1_ values fall below the minimum simulated value (≈ 0.990 vs. *>* 0.995).

Overall, our results show that linkage disequilibrium is informative for demographic inference, with different measures capturing different aspects depending on the recombination scenario. Some statistics, such as *r*^2^ and *Var*(*D*), remain robust and can be reliably used for inference, while others—like perfect positive/negative configurations or |*D*′|_1_—are more strongly affected by undetected recombination and other biological factors, limiting their direct interpretability. Importantly, these measures can be computed on real data, allowing direct comparison with theoretical expectations.

## 4 Discussion

In this study, the effects of recombination and demography are jointly investigated on different known LD measures *D*, |*D*′|, *r* and *r*^2^, as well as a newly defined set of measures we call 2-site configuration probabilities. The effect of normalization on LD has been previously studied (Kang and Rosenberg, 2019). However, we wanted to highlight the information carried by LD without normalization and the impact of *k* recombination events and demography on it. Without recombination (*k* = 0), four of the LD measures *D, r, r*^2^ and CPs are correlated and depend on the demography and the last one |*D*′| does not vary. After dissecting the different recombination types, we demonstrate that LD measures between adjacent recombination blocs (*k* = 1) are highly dependent on the type of recombination. Only incompatible recombination events affect |*D*′|. The two LD measures *D* and *r*^2^ are impacted differently by topology-changing recombination than topology-preserving recombination. Therefore, demography influences these statistics in two ways: it directly shapes the genealogical structure at each locus, affecting the LD measures themselves, and it alters the relative frequencies of recombination types, thereby modifying how recombination contributes to LD. Consequently, for adjacent recombination blocks (*k* = 1), the observed LD patterns reflect the combined effects of recombination type and demographic history. For the infinite number of recombination scenario (*k* = ∞), trees are not correlated but are affected by the same demography. In this setting, LD measures are still affected, but only through the marginal genealogical structure at each locus rather than through correlation between loci. Consequently, for long-range LD, demographic effects manifest unevenly across statistics: they have a strong impact on |*D*′|, a weaker effect on *D* and CPs, and essentially no effect on *r*^2^, which is primarily driven by sampling variance between independent genealogies.

### How can we use the different LD measures?

Analysis of both simulated and real genomic data demonstrates that different LD statistics provide complementary insights into recombination and demographic history. Specifically:

- |*D*′| is sensitive to changes in local genealogical topology caused by at least one incompatible recombination event. It can also capture correlations between gene trees induced by demographic history, especially at unlinked sites. However, empirical results show that |*D*′| may underestimate these effects when recombination events are undetected.
- *Var*(*D*), *r*, and *r*^2^ are highly similar measures of LD and robust indicators of demographic processes. While *r* and *r*^2^ are normalized, bounding their values and reducing variance near their limits ([−1, 1] for *r* and [0, 1] for *r*^2^), the variance of *D* is unbounded and more sensitive to tree topology, even for unlinked sites. These measures showed consistent patterns across diverse human populations, reflecting growth in Sub-Saharan Africans and relative decline in non-Africans.
- *E*[*r*^2^] has a predictable value for unlinked sites that is independent of demography, making it useful for detecting physical linkage (where values deviate from expectation) and for providing a baseline to identify deviations caused by other evolutionary processes.
- CPs describe the distribution of four distinct configurations between two polymorphic sites, providing finer-grained information than single-statistic-based inference alone. They can be especially useful for discriminating more complex demographic scenarios, but empirical results indicate they work best in the absence of recombination or with only a single recombination event; perfect configurations are rarely detected when recombination is effectively infinite.

Overall, all of these LD measures are informative and provide valuable tools for studying evolutionary processes, with different measures capturing distinct aspects of recombination and demography.

As shown in our study, Linkage Disequilibrium is influenced by both recombination and demography, which can be difficult to disentangle. However, by focusing on alleles that lie on the same tree, i.e., without any recombination event between them, we can isolate the effect of demography on LD. Measures computed in this context, such as *D*_0_, 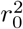, and CP_0_s, can capture evolutionary processes that shape tree topology without being affected by difference in recombination rates. This makes cross-population or cross-species comparisons more straightforward, as variation in recombination does not confound the signals. While this study focuses on demography, the same framework can be applied to other evolutionary processes such as population structure or selection. We also note that moments of the LD distribution can be analytically or computationally derived under complex evolutionary scenarios, including diploid selection, arbitrary epistasis, dominance, and variable population sizes (Ragsdale, 2022).

Recombination events may or may not alter tree topology, and even topology-changing recombinations can be difficult to detect. This limitation affects the estimation of recombination rates (Deng *et al*., 2021) and the use of recombination information in evolutionary analyses. In cases where not all recombination events are detectable, LD measures defined without recombination (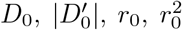, CP_0_s) cannot be directly estimated. However, most recombination events are topology-preserving, so LD measures across one recombination event (*D*_1_, *r*_1_, 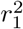, CP_1_s) closely resemble those observed without recombination. Topology-changing recombinations, though less frequent, vary with demography: demographic history not only shapes tree topology but also affects the relative frequency of recombination types. As a result, comparing LD across zero versus one recombination events (*D*_0_ vs *D*_1_) captures two sources of information: the effect of demography on tree topology and the effect of demography on the distribution of recombination types. By using detectable recombination events to segment the genome (Kerdoncuff *et al*., 2020), LD measures from different blocks can be combined strategically to capture effects from both no recombination and hidden or silent recombination events. Another possibility is to compute distributions restricted to adjacent polymorphic sites separated by likely zero or one recombination event, preserving power to detect evolutionary effects.

LD provides complementary information to SFS-based approaches. While the SFS summarizes allele frequencies across loci and is highly informative about recent demographic changes, it ignores correlations between loci. LD captures the joint distribution of alleles across loci, offering sensitivity to tree topology, recombination patterns, and subtle historical processes. Measures like CPs or *Var*(*D*) can highlight aspects of demography, such as population growth or bottlenecks, that are less apparent in the SFS, particularly when correlations between closely linked loci are informative. Nevertheless, for unlinked or effectively independent sites, the additional information provided by LD may be limited relative to the SFS.

Finally, our study demonstrates that the analytical two-locus moment approach (Hill and Weir, 1988) and that the genealogical perspective of Kingman trees and can be reconciled within a unified framework. Importantly, coalescent simulations provide a robust tool to study LD that can be directly applied to empirical data without requiring prior knowledge of the genetic map. By computing LD from phased genotypes restricted to the genomic spans defined by inferred trees, we preserve the genealogical resolution of the tree-sequence framework while mitigating biases from hidden or silent recombination. This approach allows LD statistics measured in real populations to be meaningfully compared to theoretical expectations, supporting their use for demographic inference and highlighting how different measures capture distinct aspects of population history.

## 5 Acknowledgments

GA wishes to thank the Fondation François Sommer for its financial support.

## 6 Supplementary Information I: Expectation and variance of linkage disequilibrium in a fixed tree

### 6.1 Introduction and notation

Let *T* be a (not necessarily binary) tree with *n* leaves labelled by {1, …, *n*} and let E be its edge set (neglecting the presence of a root edge, because mutations occurring on a root edge are not segregating). We introduce the notation ≺, where

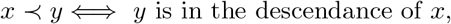

for any *x, y* may be vertices or edges of *T*.

We see *T* as the genealogy of *n* haploid individuals. Differences between the DNA sequences of these *n* individuals are due to mutations falling on *T*. We assume that each site of the sequence can be hit by **at most one mutation** (infinite-site model). Then a site that has been hit by a mutation is bi-allelic and the carriers of the mutant base form a **subtree of** *T*, **which is the set** *S*(*e*) **of leaves descending from the edge** *e* **where the mutation has fallen**:

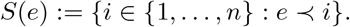

We assume that for each site hit by a mutation, the mutation falls on a random edge of E according to **the same probability distribution** *p* : E → [0, 1]. From now on, it will be convenient to think of *p*(*e*) **as the length of** *e*, as when mutations occur as homogeneous Poisson point processes on the tree. In particular, **the total length of** *T* **(sum of edge lengths) is equal to 1** and *T* is endowed with the natural distance between vertices induced by edge lengths. We also assume that **mutations occurring at different sites are independent** (conditional on *T*).

We are interested in a **pair of bi-allelic sites**, where the ancestral state is *a* for the first site, *b* for the second site and the derived/mutant state is *A* for the first site and *B* for the second site. **We let** *f*_*A*_(*T*) **(resp**. *f*_*B*_(*T*), **resp**. *f*_*AB*_(*T*)**) denote the frequency of carriers of** *A* **(resp**. *B*, **resp**. *AB***) in the leaf set of** *T*.

The **linkage disequilibrium** between these two sites is measured by the statistic

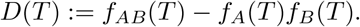

Because mutations at different sites are independent conditional on *T* and identically distributed, the expected LD is

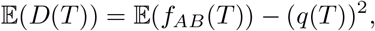

where *q*(*T*) := E(*f*_*A*_(*T*)).

For any leaf *i* = 1, …, *n*, we denote by *H*_*i*_ **the height of** *i*, or **distance between the root and** *i*, that is

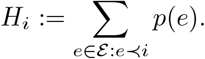

We will say that *T* is **ultrametric** when all leaves are at the same distance from the root, that is when all the *H*_*i*_’s are equal, in which case we denote their common value by *H*(*T*), **called height of** *T*.

### 6.2 Expectation of LD

#### Proposition 6.1.

*Recall that q*(*T*) *is the expected frequency of a mutation falling on T according to p. Then*

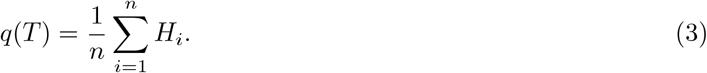

*In the ultrametric case, q*(*T*) = *H*(*T*).

*Proof*. The frequency of a mutation falling on edge *e* is #*S*(*e*)*/n*, so that

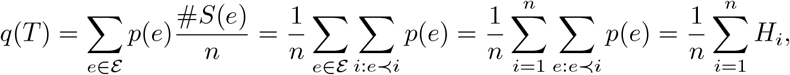

which ends the proof.

□

#### Proposition 6.2.

*Recall that f*_*AB*_(*T*) *is the frequency of the double mutant, for two mutations falling independently on T according to p. Then*

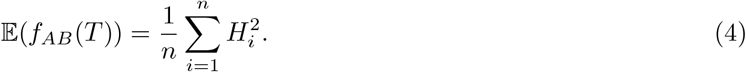

*In the ultrametric case*, E(*f*_*AB*_(*T*)) = *H*(*T*)^2^.

*Proof*. If the mutation at the first site and the mutation at the second site fall on edges *e*_1_ and *e*_2_ respectively, then

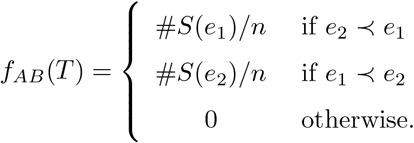

As a consequence, reasoning similarly as in the proof of the previous statement,

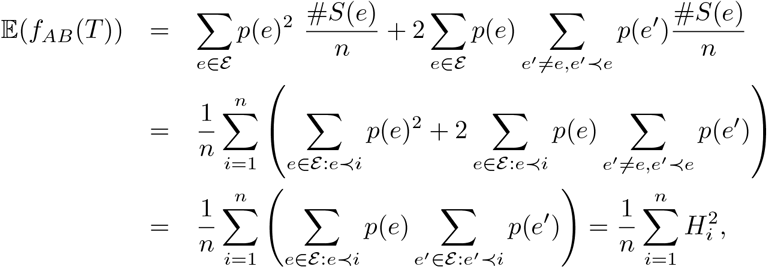

which ends the proof.

#### Corollary 6.3.

*As a consequence of the previous two propositions, the expected linkage disequilibrium in tree T is equal to*

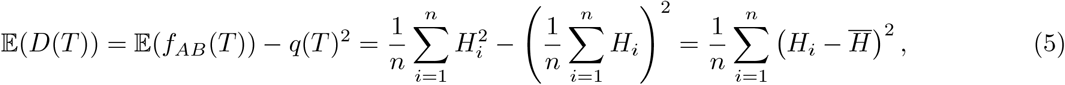

*where* 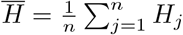 *Note that* E(*D*(*T*)) *>* 0 *except in the ultrametric case where*

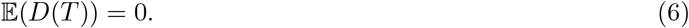

### 6.3 Variance of LD

We have to compute three quantities:

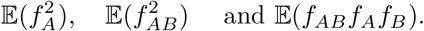

We will need to introduce, for every pair of leaves *i* and *j*, the **height** *H*_*i*∧*j*_ **of the vertex** *i* ∧ *j* **which denotes the most recent common ancestor to** *i* **and** *j*:

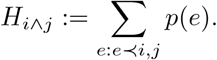

Note that

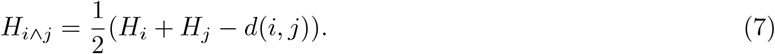

Also set

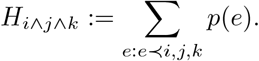

#### Lemma 6.4.

*We have*

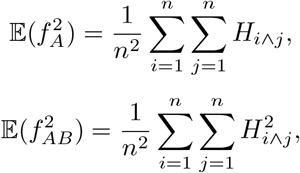

*and*

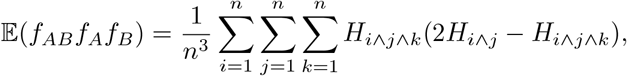

*which simplifies, in the ultrametric case, into*

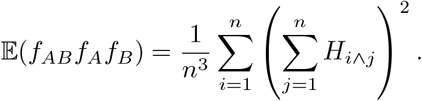

*Proof*. First,

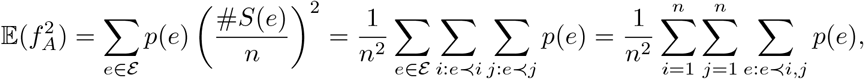

which yields the first result. Similarly,

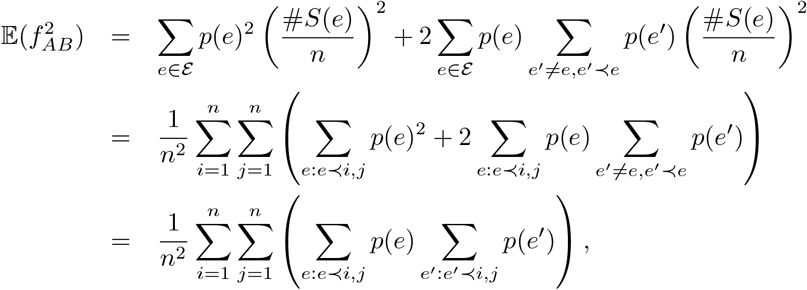

which yields the second result. Finally,

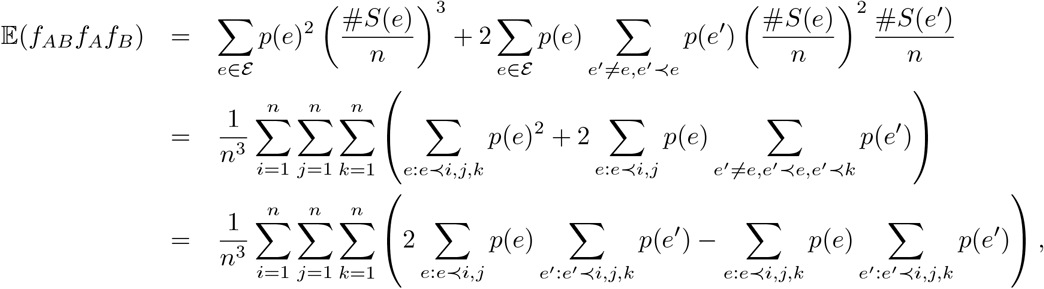

which yields

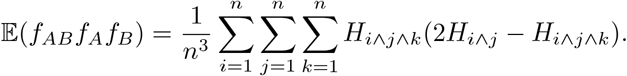

Let us try to simplify this last expression in the ultrametric case. First note that for any three leaves labelled *i, j, k*, if we set

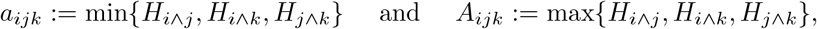

then the three quantities *H*_*i*∧*j*_, *H*_*i*∧*k*_ and *H*_*j*∧*k*_ take their values in {*a*_*ijk*_, *A*_*ijk*_}, and two of these three quantities take the value *a*_*ijk*_, which is also equal to *H*_*i*∧*j*∧*k*_. Now set

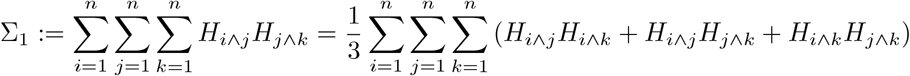

and

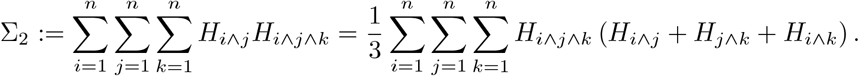

Now observe that

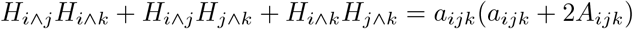

and

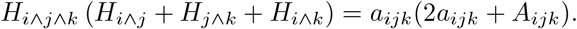

As a consequence,

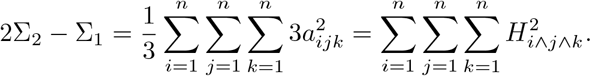

This shows that

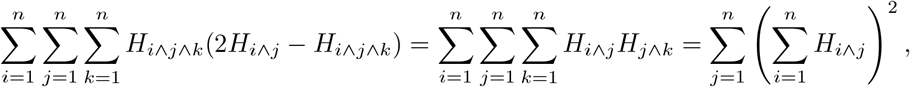

which finishes the proof.

□

We can now use the lemma to compute the second moment of *D*.

#### Proposition 6.5.

*In the general case, we have*

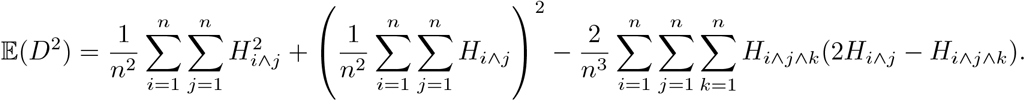

*In the case when T is ultrametric, this expression becomes (recall* E(*D*) = 0 *in this case)*

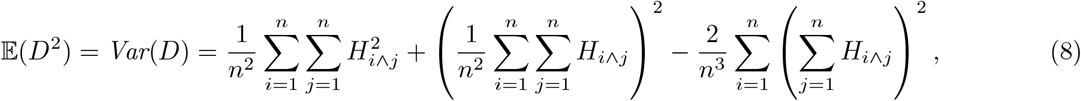

*which also reads*

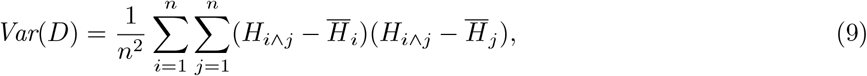

*where we have put for each i* ∈ {1, …, *n*},

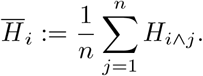

#### Remark 6.6.

*Thanks to* (7), *in the ultrametric case we have H*_*i*∧*j*_ = *H* − ^1^*d*(*i, j*). *Injecting this into* (9), *we get*

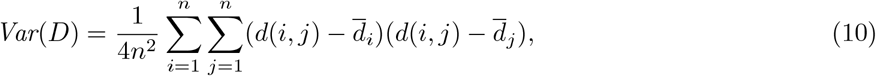

*where* 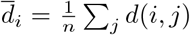. *Note that d*(*i, j*) *is equal to twice the coalescence time between leaves i and j when time is scaled so that the total length of the tree is 1. This may indicate a lead to compute the variance of D integrating over the tree T, when T is a time-inhomogeneous coalescent*.

*Proof*. By definition,

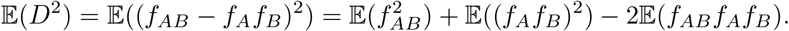

Since

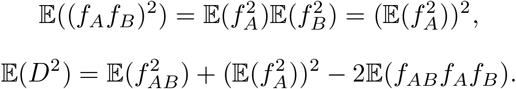

Thanks to the three expressions stated in the lemma, we get the first two expressions displayed in the proposition. Now using the notation 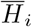, the expression in the ultrametric case reads

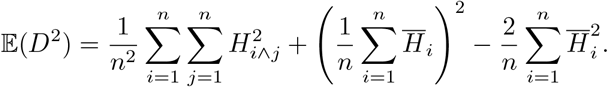

Now the second expression displayed in the statement equals

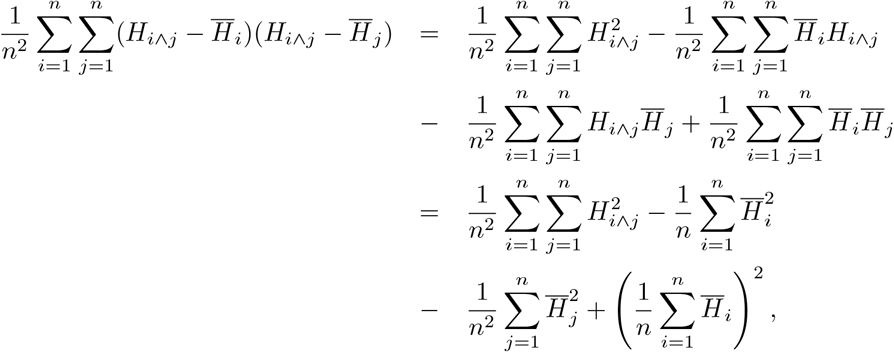

which indeed equals E(*D*^2^).

□

### 6.4 Discussion around (9)

A consequence of (9) is that 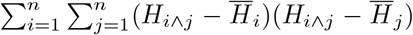 is always non-negative. This brings the question of proving independently the following general result.

#### Proposition 6.7.

*For any real, symmetric matrix M with generic element m*_*ij*_,

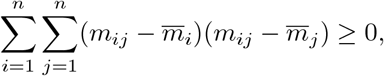

*where* 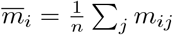.

*Proof*. Since *M* is real and symmetric, there is an orthogonal matrix *P* (^*t*^*P* = *P* ^−1^) and a diagonal matrix *D* = Diag(*λ*_1_, …, *λ*_*n*_) such that *M* = *PDP* ^−1^. Then writing *A* = *PD*, we get

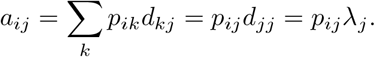

Next, since *M* = *A* ^*t*^*P*,

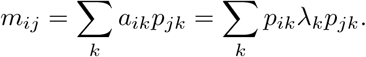

In particular,

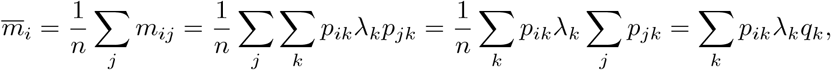

where we have set

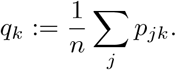

Note that Σ_*j*_ *p*_*jk*_ is the entry at row *k* of ^*t*^*Pv*, where *v* is the vector with only ones. Then

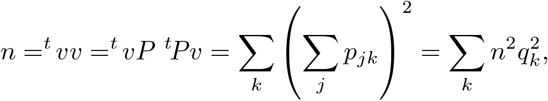

which we record as

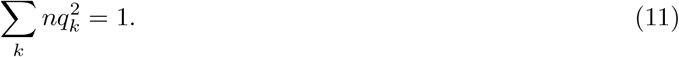

Then we have

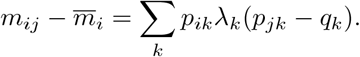

Next, defining

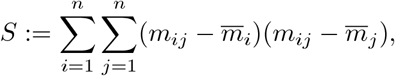

we can write

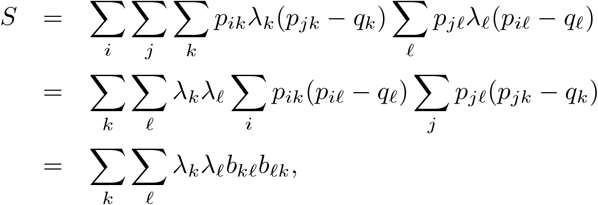

where

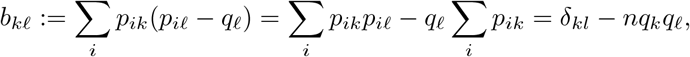

because Σ_*i*_ *p*_*ik*_*p*_*iℓ*_ is the entry (*k, ℓ*) of ^*t*^*PP* = *I*_*n*_. Finally we get

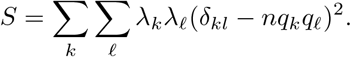

We can rewrite it as follows

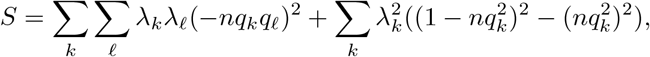

that is,

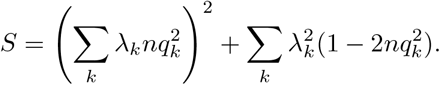

Now thanks to (11), writing 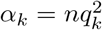, we have

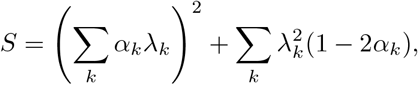

where the *α*_*k*_’s are non-negative and sum to 1. The proof ends thanks to the following result.

#### Lemma 6.8.

*For any integer n, for any non-negative* (*α*_*k*_)_*k*=1,…,*n*_ *such* that 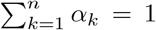 α_k_ = 1, *for any real numbers λ*_1_, …, *λ*_*n*_,

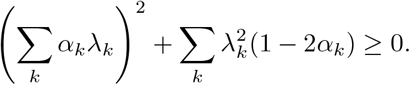

*Proof*. We reason by induction on *n*. The quantity is zero when *n* = 1. Now let *n* ≥ 2. We are going to prove that for any *α*_*k*_’s such that 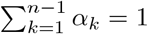, for any *t* ∈ [0, 1] and *λ*_*n*_, we have *F*_*t*_(*λ*_*n*_) ≥ 0, where

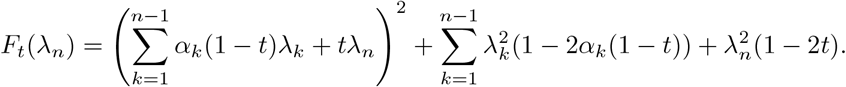

Note that *t* plays the role of *α*_*n*_ and *α*_*k*_(1 − *t*) of *α*_*k*_ for *k* ≤ *n* − 1. Also note that 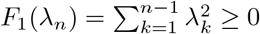, so we can assume that *t*≠ 1. Now note that *F*_*t*_(*λ*) = *a*(*t*)*λ*^2^ + 2*b*(*t*)*λ* + *c*(*t*), where

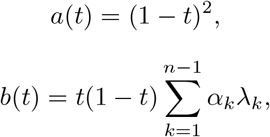

and

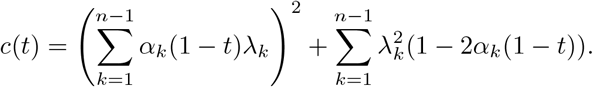

Since *t*≠ 1, *a*(*t*) *>* 0 and we only need to show that for any *t* ∈ [0, 1), *b*(*t*)^2^ − *a*(*t*)*c*(*t*) ≤ 0. Writing 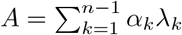, we have

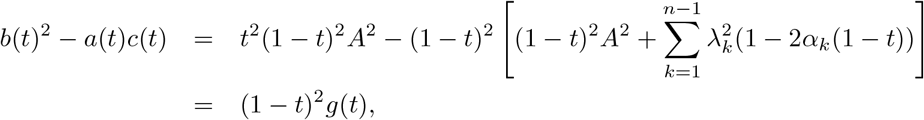

where

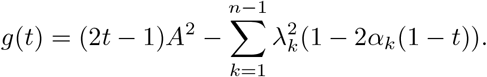

Then we only need to show that *g*(*t*) ≤ 0 for all *t* ∈ [0, 1). Since *g* is affine, it is sufficient to show that *g*(0) ≤ 0 and *g*(1) ≤ 0. Now

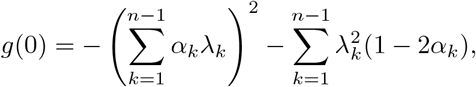

which is nonpositive by the induction hypothesis. Finally,

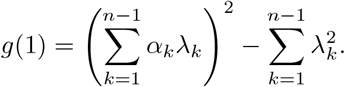

By Jensen’s inequality,

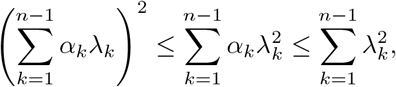

which shows that *g*(1) ≤ 0 and terminates the proof.

## 7 Supplementary Information II: Theoretical predictions for two-locus moments under a sudden demographic change in the absence of recombination

In this section, we are studying the impact of sudden change in population size *N* on two-locus moments, with a particular interest on 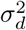, defined as:

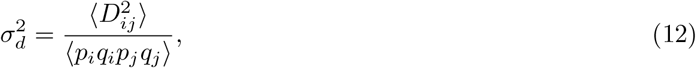

where ⟨*X*⟩ stands for the expectation of moment *X* at steady-state, *D*_*ij*_ is the linkage disequilibrium between loci *i* and *j*, and *p*_*i*_, *q*_*i*_ are allele frequencies at locus *i*.

Dynamics and expectation for 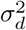 were previously obtained using a diffusion model (Ohta and Kimura, 1971) and by solving a recursion system for two-locus moments (Hill and Weir, 1988). In this section, we adopt the latter approach.

### 7.1 Theoretical framework

The usual statistics for describing linkage disequilibrium involve fourth moments and their ratios, such as *D*^2^ and *r*^2^ which are the squared covariance and squared correlation of allele frequencies respectively. It is sufficient to describe the change in the moments ⟨*p*_*i*_*q*_*i*_⟩,⟨*p*_*i*_*q*_*i*_*p*_*j*_*q*_*j*_⟩,⟨*p*_*i*_*p*_*j*_*D*_*ij*_⟩ and 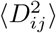 to keep track of the dynamics of all moments.

The population is assumed to be random mating with effective size *N*. It is assumed that the mutation rate per site *u* is of the order of 1*/N* or less and that the population size is sufficiently large that terms of magnitude 1*/N* ^2^, can be ignored relative to those of order 1*/N*, and the processes of drift and mutation each generation can be assumed to act additively with respect to each other.

We can describe the changes of these four moments solving:

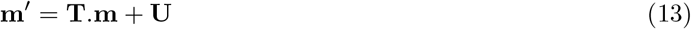

where **m** is the vector (⟨*p*_*i*_*q*_*i*_⟩,⟨*p*_*i*_*q*_*i*_*p*_*j*_*q*_*j*_⟩,⟨*p*_*i*_*p*_*j*_*D*_*ij*_⟩, 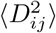), the matrix **T** is given by:

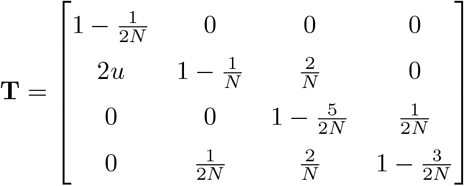

while **U** = (*u*, 0, 0, 0). **T** is similar to **I** − (**K**^*\**^ − **M**^*\**^)*/*2*N* from Equation 2 in (Hill and Weir, 1988).

Solving the equation 13 at equilibrium:

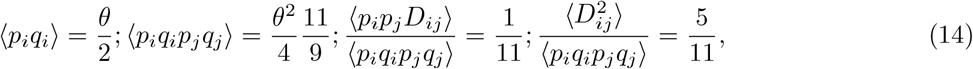

with *θ* = 4*Nu*. These results are similar to previous estimates if the recombination rate is fixed to zero (Ohta and Kimura, 1971; Hill and Weir, 1988).

The dynamics of the system are given by:

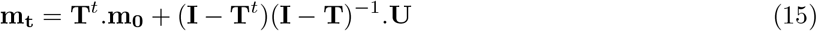

where **m**_**0**_ and **m**_**t**_ are the vector **m** at times 0 and *t*, and **I** is the identity matrix.

**Continuous time** To express our results in continuous times we use time in *N* generations (coalescent time).

We define the matrix **A** as **A** = (**I** − **T**).*N* :

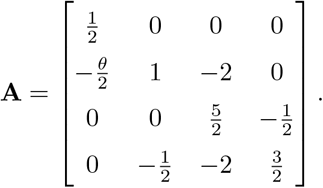

Writing Eq 15 with **A** and 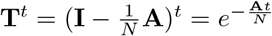, we obtain:

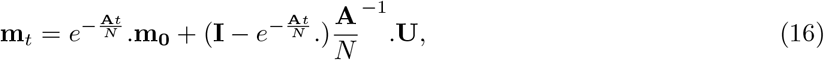

with *t*′ = *Nt* and **V** = *N*.**U**, we obtain:

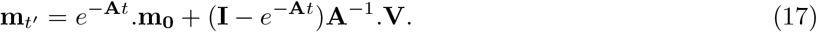

### 7.2 Demographic change

We now want to introduce a sudden change in population size. To be consistent with the demographic scenario modelled in the coalescent framework, we express time in *N*_0_ generations (size at present time): *t*′ = *N*_0_*t*. Going back in time, before the change in population size, the population was of size *N*_∞_, with *κ* the strength of the contraction and *N*_∞_ = *κ*.*N*_0_.

We consider **m**_∞_ the system at equilibrium with a population of size *N*_∞_. We express the system in *N*_0_ generations:

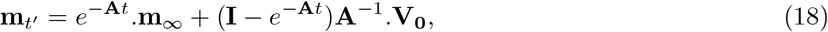

with **V**_**0**_ = *N*_0_.**U** and 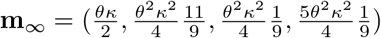

**m**_∞_ was obtained solving the equation 13 at equilibrium with *N*_∞_ = *κN* and **V**_∞_ = *κN*.**U**.

In the following, equation 18 is solved numerically, for fixed *τ* and *κ*, using MATLAB. We present 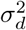 for *τ* ∈ [0.1, 0.5, 1, 10] and varying *κ* ∈ [0.01 : 100] (Figure 11). We note that *τ* estimate with the two-locus moments differ from the *τ* used in coalescent simulations by a factor two, due to the fact that coalescent simulations are based on haploid while two-locus moments are considering diploid individuals.

## 8 Supplementary Information III: LD measures estimated from Minor Allele Frequency (MAF)

## 9 Supplementary Information III: Variance of LD measures for the three recombination scenarios

## 10 Supplementary Information IV: Empirical Measurement of LD from Inferred Tree Sequences and Phased Genotypes in the 1000 Genomes Phase 3 (1KG3) Dataset, Comparing Tree-Based and Genotype-Based Approaches

### 10.1 Methods

We investigated patterns of LD across the 1KG3 (The 1000 Genomes Project Consortium, 2015) tree sequences, which were previously inferred using `tsinfer` (Kelleher *et al*., 2019). To estimate LD from empirical data, we employed two complementary approaches:

1. **Tree-based approach**: LD was computed directly from the inferred tree sequences, using branch lengths and topologies (described in more details in the following subsection).
2. **Genotype-based approach**: LD was computed from phased genotype data in VCF files. For this approach, we averaged pairwise LD statistics over all biallelic sites within the genomic span associated with each inferred tree.

To determine which method provides more reliable estimates, we compared both approaches against empirical tree sequences and simulations. Simulations were generated using `msprime` (Baumdicker *et al*., 2021), with 100 replicate datasets of 10 haploid genomes each, under a standard three-population Out-of-Africa demographic model (Gutenkunst *et al*., 2009). This comparison allowed us to assess the accuracy, robustness, and potential biases of tree-based and genotype-based LD estimates, guiding our choice of the most appropriate method for downstream analyses.

#### 10.1.1 Calculation of LD statistics

We calculate LD statistics using two complementary approaches: a *tree-based* method and a *genotype-based* (VCF-based) method.

##### Tree-based method

In the tree-based approach, we compute the expected LD directly from the genealogical trees, rather than averaging over LD between pairs of observed mutations. For each tree in the inferred tree sequence, we calculate the expected LD between pairs of branches using the tree topology and branch lengths.

As a toy example, consider the genealogy in Fig. 18. In this example, branch a is ancestral to branch b, so a pair of mutations occurring on these branches will be in positive LD. Conversely, branches a and c are either side of the root node; mutations on these branches will never be observed together on a haplotype.

**Figure 18:**
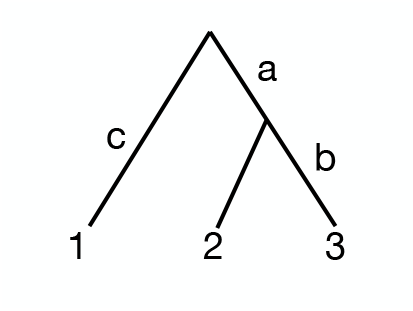
A toy tree describing the ancestry at a locus for samples 1, 2 and 3.

**Figure 19:**
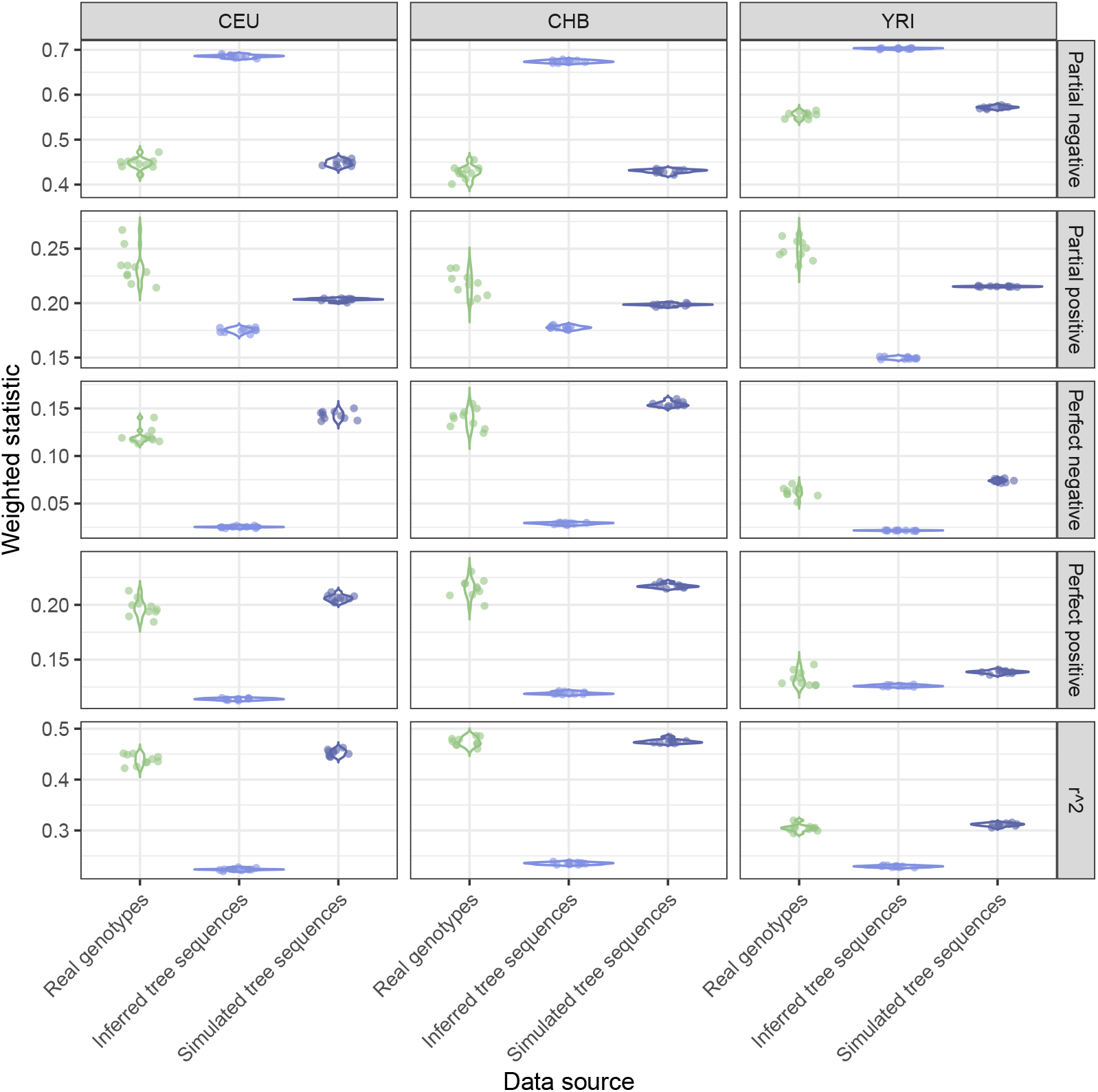
Comparison of tree-based and genotype-based LD measures within every 100th tree (containing more than two mutations) across chromosome 1 for 10 sample sets of 10 haploid genomes. We compared LD from empirical tree sequences (Kelleher *et al*., 2019) to LD from 100 simulation replicates of 10 haploid genomes, generated using msprime under a standard three-population Out-of-Africa demographic model (Gutenkunst *et al*., 2009). The LD obtained from the empirical VCF is most similar to that from the simulated trees, indicating that it is likely some form of bias resulting from tree inference which causes the discrepancy between the empirical tree-based and VCF-based measurements of LD.

Let *p*_*a*_ denote the allele frequency of a mutation occurring on branch *a*, given by the proportion of samples descending from *ba*, and the same for *p*_*b*_ and *p*_*c*_. In Fig. 18, *p*_*a*_ = 2*/*3, *p*_*b*_ = 1*/*3 and *p*_*c*_ = 1*/*3. Conditional upon this topology, we know that *p*_*ab*_ = *p*_*b*_, so the expected *r*^2^ between a mutation on branch *a* and a mutation on branch *b* is

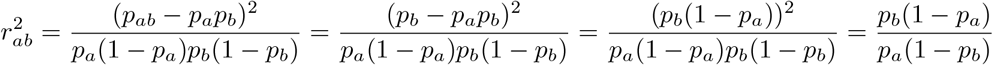

Therefore,

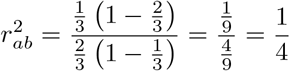

Instead, for mutations *a* and *c*, we have that

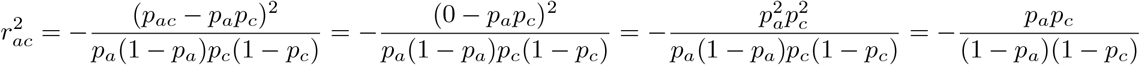

With *p*_*c*_ = 1 − *p*_*a*_ leads to 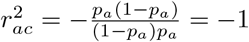

Other LD statistics can be derived using the same approach. This framework naturally extends to mutations occurring on different trees. Once the LD values associated with a given topology are determined, the remaining step is to weight them by the probability of observing the corresponding mutations.

Under the infinite-sites model, the probability of a mutation occurring on branch *b* is *l*_*b*_*/L*, where *l*_*b*_ is the branch length and *L* is the total branch length of the tree. Averaging 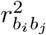 over all pairs (*b*_*i*_, *b*_*j*_), with the weight of each branch as 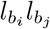 in the tree yields the tree-wide expected LD — i.e. the expected distribution of an LD statistic at that locus.

##### Genotype-based method

We also calculate LD statistics directly from the variant calls in the VCF files. In this approach, we average over observed pairs of mutations within a genomic segment, where a segment is defined as the span of a single inferred tree in the tree sequence.

For both methods, LD measures were computed at three recombination scales:

- *D*_0_: LD within a single tree, averaged across every 100th tree on all autosomes.
- *D*_1_: LD between neighboring trees, averaged across every 100th tree on all autosomes.
- *D*_∞_: Approximate LD at an infinite recombination distance, estimated by randomly sampling 100 tree pairs from chromosome pairs (1/2, 3/4, …, 21/22).

For each tree pair, we calculated both tree-based and genotype-based LD statistics. The genotype-based measurements used the genomic span of the tree but relied on genotypes recorded in the 1KG3 VCF. We restricted the analysis to biallelic sites and only considered comparisons where both trees contained more than two sites. We also eliminated trees which had a span greater than 1e5 base pairs.

#### 10.1.2 Dataset

To construct our dataset, we created 10 sample sets of 10 haploid genomes for each 1KG3 population, grouped by super-population: Sub-Saharan Africa (AFR) – Yoruba in Ibadan, Nigeria (YRI), Luhya in Webuye, Kenya (LWK), Esan in Nigeria (ESN), Mende in Sierra Leone (MSL), Gambian in Western Divisions (GWD); Americas (AMR) – Colombians in Medellin, Colombia (CLM), Peruvians in Lima, Peru (PEL), Puerto Ricans in Puerto Rico (PUR), Mexican Ancestry in Los Angeles, USA (MXL); East Asia (EAS) – Han Chinese in Beijing (CHB), Southern Han Chinese (CHS), Japanese in Tokyo (JPT), Kinh in Ho Chi Minh City, Vietnam (KHV); Europe (EUR) – Utah Residents with Northern and Western European Ancestry (CEU), Finnish in Finland (FIN), British in England and Scotland (GBR), Toscani in Italy (TSI), Iberian populations in Spain (IBS); and South Asia (SAS) – Gujarati Indians in Houston, Texas, USA (GIH), Punjabi in Lahore, Pakistan (PJL), Bengali in Bangladesh (BEB), Sri Lankan Tamil in the UK (STU), Indian Telugu in the UK (ITU). We removed individuals with a child, parent, or grandparent in the dataset to minimize relatedness and separated each individual’s two haploid genomes into different sample sets to approximate a haploid sample. While all populations were used in the overall analyses, for direct comparison with simulations we focused on three representative populations: CEU, CHB, and YRI.

#### 10.1.3 msprime simulations

In order to understand the extent to which the LD of simulated tree sequences compares to that obtained empirically, we conducted `msprime` simulations under a three-population Out of Africa model from Gutenkunst *et al*. (2009). This demographic model describes an ancestral human population in Africa, and a subsequent European-Asian population split with migration and population growth. Model parameters reflecting the maximum likelihood values for CHB, CEU and YRI, as recorded in the `stdpopsim` library (Lauterbur *et al*., 2023). We constructed sample sets again by selecting 10 haploid genomes from each population (i.e. avoiding that both copies of one individual’s genome be included in the same sample set).

### 10.2 Discrepancy of tree and genotype-based LD measurement in the 1KG3 data

We found that nearly all LD statistics differed between tree-based and genotype-based measurements. To understand these discrepancies, we compared LD computed directly from tree sequences to LD derived from simulated mutations placed on the same trees. We applied this approach both to empirically inferred tree sequences and to tree sequences simulated under a three-population Out-of-Africa model (Gutenkunst *et al*., 2009). In both cases, tree-based LD closely matched the LD measured from simulated mutations (Figures 19). However, tree-based LD deviated significantly from LD measured directly from empirical genotype data. This suggests that errors in tree inference distort genealogical structure in ways that affect associations between sites. In contrast, SNP-based LD closely matched LD from trees simulated under the Out-of-Africa model, suggesting that this model captures real LD patterns well.

Therefore, in the main text, we show LD obtained from genotypes, as described above. These findings indicate that current methods for tree inference do not fully preserve the empirical patterns of LD observed in genetic variation. Improving tree reconstruction methods—or selectively analyzing well-resolved regions of inferred trees—may be necessary to make genealogical LD measurements a more reliable alternative to traditional genotype-based LD calculations.

